# Single-cell RNA sequencing-based characterization of resident lung mesenchymal stromal cells in bronchopulmonary dysplasia

**DOI:** 10.1101/2021.06.18.448928

**Authors:** I. Mižíková, F. Lesage, C. Cyr-Depauw, D. P. Cook, M. Hurskainen, S.M. Hänninen, A. Vadivel, P. Bardin, S. Zhong, O. Carpen, B. C. Vanderhyden, B. Thébaud

**Affiliations:** Sinclair Centre for Regenerative Medicine, Ottawa Hospital Research Institute, Ottawa, ON, Canada; Department of Cellular and Molecular Medicine, University of Ottawa, Ottawa, Ontario, Canada; Cancer Therapeutics Program, Ottawa Hospital Research Institute, Ottawa, Ontario, Canada; Division of Pediatric Cardiology, New Children’s Hospital, Helsinki University Hospital and University of Helsinki, Helsinki, Finland; Pediatric Research Center, New Children’s Hospital, University of Helsinki and Helsinki University Hospital, Helsinki, Finland; Precision Cancer Pathology, Department of Pathology and Research Program in Systems Oncology, University of Helsinki and HUS Diagnostic Center, Helsinki University Hospital, Helsinki, Finland; Department of Pediatrics, Children’s Hospital of Eastern Ontario (CHEO) and CHEO Research Institute, University of Ottawa, Ottawa, Ontario, Canada; Department of Obstetrics and Gynecology, University of Ottawa/The Ottawa Hospital, Ottawa, Ontario, Canada

## Abstract

Late lung development is a period of alveolar and microvascular formation, which is pivotal in ensuring sufficient and effective gas exchange. Defects in late lung development manifest in premature infants as a chronic lung disease named bronchopulmonary dysplasia (BPD). Numerous studies demonstrated the therapeutic properties of exogenous bone marrow and umbilical cord-derived mesenchymal stromal cells (MSCs) in experimental BPD. However, very little is known regarding the regenerative capacity of resident lung MSCs (L-MSCs) during normal development and in BPD. In this study we aimed to characterize the L-MSC population in homeostasis and upon injury. We used single-cell RNA sequencing (scRNA-seq) to profile *in situ Ly6a*^+^ L-MSCs in the lungs of normal and O_2_-exposed neonatal mice (a well-established model to mimic BPD) at three developmental timepoints (postnatal days 3, 7 and 14). Hyperoxia exposure increased the number, and altered the expression profile of L-MSCs, particularly by increasing the expression of multiple pro-inflammatory, pro-fibrotic, and anti-angiogenic genes. In order to identify potential changes induced in the L-MSCs transcriptome by storage and culture, we profiled 15,000 *Ly6a*^+^ L-MSCs after *in vitro* culture. We observed great differences in expression profiles of *in situ* and cultured L-MSCs, particularly those derived from healthy lungs. Additionally, we have identified the location of L-MSCs in the developing lung and propose *Serpinf1* as a novel, culture-stable marker of L-MSCs. Finally, cell communication analysis suggests inflammatory signals from immune and endothelial cells as main drivers of hyperoxia-induced changes in L-MSCs transcriptome.

## 1. INTRODUCTION

Late lung development represents an important period in lung maturation marked by an exponential increase in the gas exchange surface area by forming the most distal respiratory units, the alveoli. Within these units, respiration takes place across a thin (0.2 - 2μm) alveolo-capillary barrier. Formation of alveolar structures, a process known as alveolarization, is facilitated by spatially and temporarily coordinated interactions between diverse cell types and the pulmonary microenvironment [1]. Defects in late lung development in humans manifest as bronchopulmonary dysplasia (BPD), a multifactorial disease occurring as a consequence of premature birth, respiratory distress, and associated treatments in neonatal intensive care. BPD is the most common chronic disease in children and a leading cause of death in children under the age of 5 [1,2]. BPD is also associated with neurodevelopmental delay, increased incidence of asthma, re-hospitalizations and early-onset emphysema [3,4].

To date, multiple studies have demonstrated the lung protective effects of exogenous, bone marrow (BM)- or umbilical cord (UC)-derived, mesenchymal stromal cells (MSCs) in experimental BPD models [5–10]. The discovery of lung resident (L-)MSCs prompted questions regarding the apparent insufficient regenerative capacity of L-MSCs in lung injury [11]. Characterizing the L-MSC population in homeostasis and upon injury is pivotal in understanding the apparent contradiction between the therapeutic effects of exogenous MSCs, while the resident population fails to prevent neonatal lung injury from occurring. However, very little is currently known about the role of L-MSCs in postnatal lung development and in BPD. Lung stromal cells, including lipofibroblasts, myofibroblasts and matrix fibroblasts are a potent source of inter-cellular signaling and are known to play an important role in BPD pathogenesis [12]. However, how L-MSCs communicate with other cell populations and contribute to the development of BPD remains unknown.

While most authors report that L-MSCs can differentiate, to some extent, into chondroblasts, osteoblasts and adipocytes [13], form colonies *in vitro* [13,14], and express classical MSC markers THY1 (CD90), NT5E (CD73) and ENG (CD105) [13,15], no L-MSC-specific marker has yet been established. Due to the lack of standardization for L-MSC identification, as well as differences in expression profiles between species, no single marker has been broadly accepted. Lung mesenchymal progenitor cell markers have been proposed [13,15–18], including LY6A, often referred to as SCA-1 (Stem cell antigen 1) [16–19]. LY6A was proposed as a defining progenitor marker for mesenchymal cell lineages in the lung [19] and LY6A^+^ mesenchymal lung cells were shown to promote colony formation, proliferation and differentiation of epithelial progenitor cells [20].

In the study presented here we identify, for the first time, the transcriptome of *Ly6a*^+^ L-MSCs in heathy and diseased developing mouse lungs. We hypothesized, that O_2_-exposure (a well-established model to mimic BPD) significantly impacts the phenotype and function of L-MSCs, as well as cellular communication between L-MSCs and other cell populations in the developing lung. We identify perturbations to the phenotype and functional properties of L-MSCs in this model. Furthermore, we report extensive single-cell RNA sequencing (scRNA-seq) profiling of L-MSCs in the lungs of 36 healthy and O_2_-exposed mice at three developmental timepoints (P3, P7, and P14). Finally, we investigate cultured *Ly6a*^+^ L-MSCs and *Ly6a*^−^ mouse lung stromal cells by scRNA-seq. We identify changes in L-MSCs transcription profile induced by storage and culture and present novel, culture-stable marker for this rare progenitor population.

## 2. MATERIALS AND METHODS

### 2.1 Experimental animals

Pregnant C57BL/6N mice were purchased from Charles Rivers Laboratories, Saint Constant, QC, Canada at embryonic day (E)15. Mice were housed by the Animal Care and Veterinary Service of the University of Ottawa in accordance with institutional guidelines. All study protocols were approved by the animal ethics and research committee of the University of Ottawa (protocol OHRI-1696) and conducted according to guidelines from the Canadian Council on Animal Care (CCAC). Mouse pups born on the same day, were randomized at the day of birth [postnatal day (P)0] and divided into equal-sized litters of 6-8 pups/cage. Cages were then maintained either in room air (normoxia, 21% O_2_), or in normobaric hyperoxia (85% O_2_) until the day of harvest. The hyperoxic environment was maintained in sealed plexiglass chambers with continuous oxygen monitoring (BioSpherix, Redfield, NY). Mice were maintained in 12/12 hours light/dark cycle and received food ad libidum. In order to avoid confounding factors associated with oxygen toxicity, nursing dams were rotated between normoxic and hyperoxic group every 48 hours. Euthanasia was performed by an intraperitoneal (i.p.) injection of 10 μl/g Pentobarbital Sodium (CDMV, Saint-Hyacinthe, QC, Canada).

### 2.2 Lung isolation

Mouse pups designated for mean linear intercept (MLI) assessment or fluorescent in situ hybridization (FISH) were euthanized at P7 and P14, respectively. Following euthanasia, the chest was opened, mice were tracheotomized and lungs were installation-fixed for 5 minutes at 20cm H2O hydrostatic pressure. Lungs designated for histological assessment were fixed with 1.5% (w/v) paraformaldehyde (PFA) (Sigma-Aldrich, Oakville, ON, Canada) and 1.5% (w/v) glutaraldehyde (Sigma-Aldrich, Oakville, ON, Canada) in 150mM HEPES (Sigma-Aldrich, Oakville, ON, Canada). Lungs designated for FISH were fixed with 4% (w/v) PFA (Sigma-Aldrich, Oakville, ON, Canada). In both instances, lungs were kept in the fixation solution for 48 hours at 4°C and collected for embedding in paraffin. Paraffin-embedded tissue blocks designated for histological analyses were sectioned at 3 or 4μm as needed. Tissue dehydration, paraffin embedding and sectioning were performed by the University of Ottawa Louise Pelletier Histology Core Facility.

Mouse pups designated for lung cells isolation and fluorescence activated cell sorting (FACS) analyses were euthanized at P7. Mice also received an i.p. injection of 10 mU/g Heparin Sodium (LEO Pharma INc., Thornhill, ON, Canada). Following euthanasia, the chest was opened and the left atrium was perforated. Lungs were perfused through the right ventricle with 5 ml of 25 U/ml Heparin Sodium in DPBS supplemented with Mg^2+^/Ca^2+^ (ThermoFisher Scientific, Burlington, ON, Canada) until white. Lungs were removed from the thoracic cavity, dissected into individual lobes, and digested in enzyme mix at 37°C by gentleMACS™ Octo Dissociator (Miltenyi Biotech, Bergisch Gladbach, Germany). The detailed procedure, as well as enzyme mixture contents are provided in Supplementary Method S1. The suspension was then centrifuged and the resulting pellet was washed with 5 ml of 5% FBS (Sigma-Aldrich, Oakville, ON, Canada) in 1× DPBS (Lonza, Basel, Switzerland), filtered through 70 μm filter (Corning Life Sciences, Tewksbury, MA, USA) and centrifuged again. The resulting pellet was resuspended in 1ml of cold RBC lysis buffer (ThermoFisher Scientific, Burlington, ON, Canada) for 3 minutes at room temperature (RT). The cell suspension was then diluted with 5ml of 5% FBS solution, centrifuged and washed twice.

A detailed flowchart illustrating the allocation of each mice to respective experimental groups is depicted in Supplementary figure 1.

### 2.3 Mean linear intercept (MLI) measurement

Paraffin-embedded tissue blocks were sectioned at 4μm, stained with hematoxylin and eosin (H&E) stain, and scanned using the Axio Scan.Z1 (Zeiss, Oberkochen, Germany). The mean linear intercept (MLI) was estimated with Fiji/ImageJ software using a 64-point grid as described previously [21]. A total of 20 randomly selected 500μm×500μm fields of view were assessed in each lung.

### 2.4 Fluorescent activated cell sorting (FACS)

The number of cells in the single-cell suspension was estimated using the EVE NanoEnTek automatic cell counter and a total of 1×106 cells/sample were resuspended in 550 μl of FACS buffer (5% (v/v) FBS and 1mM EDTA in 1×DPBS). Cells were then incubated at RT in the dark with 2 μl/1×10^6^ cells of CD16/32 antibody for 15 minutes. Following blocking, cells were centrifuged and pellets were resuspended in 1:100 mixture of panel of antibodies: FITC-CD31, AF647-CD45, Pe/Cy7-CD326, and BV421-LY-6A/E (Supplementary table 1). Cells were incubated with antibodies for 20 minutes in dark at RT, pelleted and washed 3x with FACS buffer. FACS was performed immediately using a MoFlo XDP (XDP, Beckman Coulter, Fullerton, CA, USA) and compensation and analysis was done using Summit v.5.4 at the Ottawa Hospital Research Institute (OHRI) StemCore facility.

### 2.5 Cell culture and storage

The detailed procedure is provided in Supplementary Method S2.

### 2.6 Colony formation assay

The detailed procedure is provided in Supplementary Method S3.

### 2.7 MSCs surface marker profiling

Cultured, passage 3 CD31^−^/CD45^−^/EpCAM^−^/LY6A^+^ L-MSCs were profiled for MSC surface markers by flow cytometry. Briefly, 3×10^5^ cells/sample were resuspended in 200 μl of FACS buffer in 96-well plate and incubated at RT in the dark with 2 μl/1×10^6^ cells of CD16/32 antibody for 15 minutes. Cells were then divided to 3 equal fractions, centrifuged and resuspended in one of the following 1:100 mixture of antibodies: i) BV421-CD31, Pe/Cy7-conjugated CD326, PE-CD73, and AF488-D105; ii) AF647-CD45, BV421-LY-6A/E, PE-conjugated CD34, and AF488-CD146; iii) PB-CD90.2 (Supplementary table 1). Cells were incubated with antibodies for 20 minutes in dark at RT, pelleted and washed 3x with FACS buffer. Flow cytometry was performed immediately using a MoFlo XDP (XDP, Beckman Coulter, Fullerton, CA, USA) and compensation and analysis was done using Summit v.5.4 at the OHRI core facility.

### 2.8 Osteogenic differentiation

The detailed procedure is provided in Supplementary Method S4.

### 2.9 Adipogenic differentiation

The detailed procedure is provided in Supplementary Method S5.

### 2.10 Chondrogenic differentiation

The detailed procedure is provided in Supplementary Method S6.

### 2.11. Fluorescent in situ hybridization

The detailed procedure, as well as a list of used probes are provided in Supplementary Method S7.

### 2.12. Multiplexing samples for scRNA-seq

Multiplexing was performed according to the MULTI-seq protocol [22]. The detailed procedure is provided in Supplementary Method S8.

### 2.13. scRNA-seq library preparation and sequencing

Single-cell suspensions were processed using the 10x Genomics Single Cell 3’ v3 RNA-seq kit by Ottawa Hospital Research Institute Stem Core Laboratories. Gene expression libraries were prepared according to the manufacturer’s protocol. MULTI-seq barcode libraries were retrieved from the samples and libraries were prepared independently, as described previously[22]. Final libraries were sequenced on the NextSeq500 platform (Illumina) to reach an approximate depth of 20,000-25,000 reads/cell.

### 2.14. scRNA-seq data analyses and quantification

#### Processing and demultiplexing

Raw sequencing reads were processed using CellRanger v3.0.2 for lung homogenate sample and v3.1.0 for cultured cells, aligning reads to the mm10 build of the mouse genome. Except for explicitly setting --expect-cells=25000, default parameters were used for all samples. MULTI-seq barcode libraries were trimmed prior to demultiplexing to 28bp using Trimmomatic (v0.36). Demultiplexing was performed using the deMULTIplex R package (v1.0.2) as described previously[22,23]. Only cells positive for a single barcode were kept for further analysis and sample annotations were added to all cells in the data set.

#### Quality control, integration, and clustering

All main processing steps were performed with Seurat v.4.0.0[24]. Quality control was performed independently on each library to find appropriate filtering thresholds. Expression matrices were loaded as Seurat objects into R. Only cells with > 200 genes detected and < 20% of UMIs mapped to mitochondrial genes were retained. Each unique sample was split based on MULTI-seq sample barcodes into a separate Seurat object. SCTransform[25] was used to normalize samples, select highly variable genes, and to regress out cell cycle and cell stress effects. To eliminate batch effects or biological variability effects on clustering, the data integration method implemented by Seurat for SCTransform-normalized data was performed, using the SelectIntegrationFeatures(), PrepSCTIntegration(), FindIntegrationAnchors(), and IntegrateData() functions. PCA was performed on the top 3000 variable genes and the data was clustered at a low resolution (dims=1:30, resolution=0.2 for lung homogenate data and 0.1 for cultured MSCs) with the Louvain algorithm implemented in the FindClusters() function in Seurat. Cell populations were identified with a simple Wilcoxon rank sum test with the FindAllMarkers() function in Seurat.

In the case of stromal cells from lung homogenates, a previously published, publicly available scRNA-seq dataset from newborn mice was re-analyzed[23]. A novel Ly6a^+^ L-MSC population was identified based on the expression of Ly6a. New cell type labels for stromal populations were then added to the Seurat object containing all data.

#### Differential expression analysis (DSA), gene set enrichment analysis (GSEA) and functional enrichment analysis

To identify differentially expressed genes in response to hyperoxia or as a result of mouse age, we used the R package muscat (v1.4.0). Pseudobulk expression profiles were generated for each sample in each cluster and differential expression was tested between groups associated with the experimental conditions. Genes with an adjusted p-value < 0.05 and a detection rate ≥ 10% in at least one of the conditions tested were considered significant. To further identify gene sets associated with differentially expressed genes, we used the R package fgsea (v1.16.0). List of gene sets comprised all GO terms, KEGG pathways, Reactome pathways, and the MSigDB Hallmark gene sets acquired from the Molecular Signatures Database (v7.2)[26]. Gene sets with an adjusted p-value < 0.05 were considered significantly enriched. Normalized enrichment score (NES) was used to assess whether gene sets were associated with upregulated or downregulated genes. Functional enrichment analysis (FEA) for selected ligands produced by Ly6a^+^ L-MSCs were performed using the online Metascape tool[27]. Summary pathways relevant to lung were considered.

#### Cell communication inference

To explore cell communication networks behind the developmental age, or hyperoxia-specific effects, we utilized the R package nichenetr (v1.0.0), which uses information about expression of cognate ligands, receptors, signaling pathways, and genomic targets to infer cell communication patterns[28]. Differential gene expression analysis for P3 vs. P14, or hyperoxia vs. normoxia groups were used in the NicheNet analysis. To prioritize results, analysis was limited to signaling contributing to the effects in receiver cell types with >200 differentially expressed genes at P14 or in response to hyperoxia, but included all cell types as potential ligand senders. Background expression of genes was specified with default approach used in NicheNet’s pipeline, using all genes with >10% detection in a given cluster. While using cells from both experimental conditions, developmental age, or hyperoxia-induced ligands from cell types that increase in proportion with age or in hyperoxia samples were prioritized. For each “receiver” cell population, top 10 ligands predicted to drive developmental age, or hyperoxia-induced responses were selected based on the Pearson correlation coefficient between the ligand-target regulatory potential score of each ligand and the target indicator vector. Further, we assessed whether the expression of ligands and receptors was upregulated, or whether the populations expressing the ligands increased in proportion in P14 or hyperoxia samples, respectively. Finally, potential target genes were inferred. Summaries of ligand-receptor interactions are represented in circos plots.

### 2.15. Statistical analysis

All statistical analyses were performed with GraphPad Prism 8.0. The presence of potential statistical outliers was determined by Grubbs’ test. Data are presented as means ± SD. Differences in case of two-member groups were evaluated either by unpaired Student’s *t*-test, or multiple unpaired Student’s *t*-test with correction for multiple comparisons using the Holm-Šidák method. P values ˂ 0.05 were considered as significant and depicted as following: P values ˂ 0.05: *; P values ˂ 0.01: **; P values ˂ 0.001: ***; P values ˂ 0.0001: ****.

## 3. RESULTS

### 3.1 The developing murine lung contains a population of L-MSC marked by the expression of *Ly6a*

In order to understand the expression patterns unique to LY6A^+^ L-MSCs in the developing lung, we took advantage of a publicly available scRNA-seq dataset from newborn mice[23]. Within this dataset, we analyzed 7,994 stromal cells from normoxia or hyperoxia-exposed developing mouse pups on postnatal days (P)3, 7, and 14, clustered into 6 distinct populations (Fig. 1A). Based on the expression pattern of commonly used MSC markers (Fig. 1B, Supplementary fig. 2A) we selected *Ly6a* as most suitable marker to identify L-MSC in lung stroma. We then subsetted the *Ly6a*^+^ cells, belonging almost exclusively to the *Col14a1*+ fibroblasts, as a separate, seventh cluster (Fig. 1B). Differential gene expression analysis revealed that *Ly6a*^+^ L-MSCs could be characterized by the expression of additional markers, including *Lum*, *Serpinf1*, or *Dcn*, with *Lum* being the single most unique identifier of the population (Fig. 1C, Supplementary table 2). It was previously shown to inhibit migration, invasion, and tube-formation in BM-MSCs[29], and was implicated in epithelial-mesenchymal transition and fibrocyte differentiation[30]. While *Ly6a*^+^ L-MSCs expressed additional MSC markers *Mcam*, *Alcam* and *Eng*, their expression did not serve as a reliable indicator of *Ly6a*^+^ L-MSCs (Fig. 1D).

**Figure 1.**
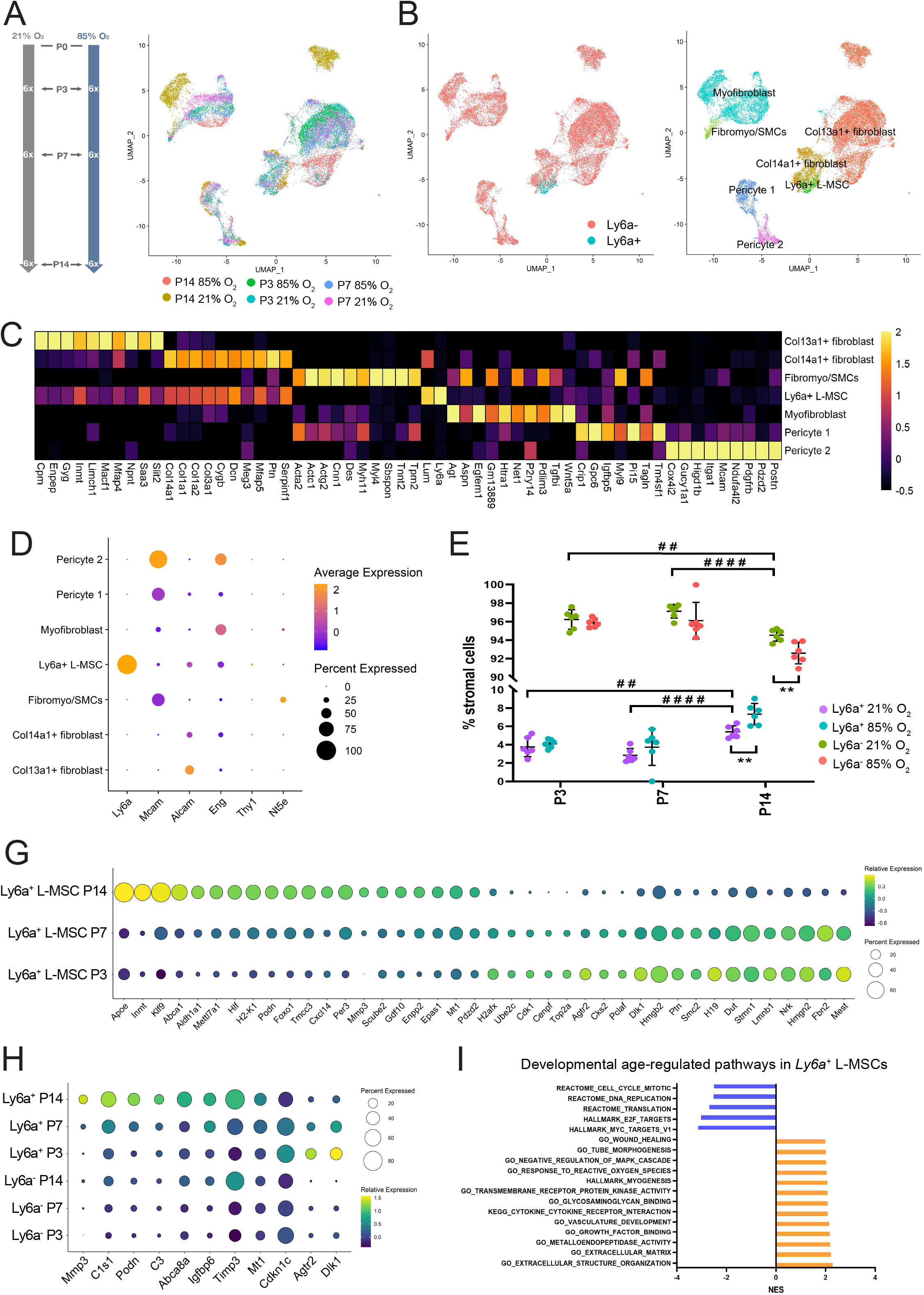
Gene expression profile of *Ly6a*^+^ L-MSCs during late lung development. **(A)** Six clusters of stromal cells were previously identified in developing lungs. In the dataset re-analyzed here mice were exposed to room air (21% O_2_) or hyperoxia (85%O_2_) from P1 onwards and lungs were harvested at P3, P7 and P14. UMAP plot depicting the distribution of lung stromal cells based on the developmental age and oxygen exposure. **(B)** UMAP plots showing the expression of *Ly6a* mRNA (left panel) within the lung stroma and new cluster identities, including the *Ly6a^+^* L-MSCs. **(C)** Heatmap of top ten most differentially expressed genes across stromal clusters depicted in panel (B). **(D)** Dotplot depicting expression of routine MSC markers in lung stromal populations. **(E)** Relative contribution of *Ly6a*^+^ and *Ly6a*^−^ cells in developing lung stroma at P3, P7 and P14. n = 6 animals/group. Data are presented as means ± SD. Statistical analyses were performed with GraphPad Prism 8.0 and the presence of potential statistical outliers was determined by Grubbs’ test. Significance was evaluated by multiple unpaired Student’s *t*-test with Holm-Šidák correction for *Ly6a*^+^ and *Ly6a*^−^ cells separately. *P* values < 0.05 were considered significant and are depicted. **(F)** Dotplot depicting the expression of most differentially expressed genes in *Ly6a*^+^ L-MSCs during normal lung development. **(G)** Dotplot depicting the expression of genes that are differentially expressed specifically in *Ly6a*^+^ L-MSCs and not in other lung stromal clusters during normal lung development. **(H)** Selected developmental age-associated signalling pathways in the *Ly6a*^+^ L-MSC cluster identified by gene set enrichment analysis (GSEA). All terms are significantly enriched (adjusted p-value < 0.05). Normalized enrichment scores (NES) values were computed by gene set enrichment analysis on fold change-ranked genes. Expression values in Heatmap represent Z-score-transformed log(TP10k+1) values. Expression levels in Dotplots and UMAP plots are presented as log(TP10k+1) values. Log(TP10k+1) corresponds to log-transformed UMIs per 10k.

### 3.2 The transcription profile and signaling activity of *Ly6a*^+^ L-MSCs change significantly during postnatal lung development

We first aimed to understand how the L-MSC population changes in the postnatal developing lung. While the size of the population remained unchanged between P3 and P7, the second week of lung development in healthy mice was associated with an increase in the size of the *Ly6a*^+^ stromal population (Figure 1E, Supplementary table 3). Similarly, differential state analysis (DSA) in normally developing lungs revealed that most changes in gene expression occurred in L-MSC between P7 and P14 (Fig. 1F, Supplementary table 4). Although the expression of genes such as *Apoe*, *Inmt*, *Klf9* and *Abca1* was drastically increased in L-MSCs, these genes were also considerably upregulated in *Ly6a*^−^stromal cells (Supplementary table 4). The largest L-MSC - specific expression changes were observed for *Mmp3*, *C1s1*, *Podn*, *Dlk1*, and *Agtr2* (Fig. 1G). Gene set enrichment analysis (GSEA) identified extracellular matrix (ECM) formation, vascular development, and wound healing among the activated pathways (Fig. 1H, Supplementary table 5).

Next, to further understand how L-MSCs send and receive signals during postnatal development, we performed a cell communication analysis. We inferred developmental age-induced cellular communications between *Ly6a*^+^ L-MSCs and other lung populations using the NicheNet tool [23,28] (Fig. 2A, Supplementary fig. 3-5, Supplementary table 6). During development L-MSCs received signals from several cell populations, including endothelial cells, interstitial macrophages (Int Mf), alveolar epithelial type 2 (AT2) cells, and stromal cells (Fig. 2A). *Col4a1*, *Fat1*, *Hmgb2*, *Vcam1* and *Hc* were identified as most potent ligands, targeting numerous downstream genes in the developing L-MSCs, including *Klf9*, *Top2a* and other strongly de-regulated genes (Fig. 2A-B, Figure 1F, Supplementary table 4). Furthermore, L-MSCs produced numerous ligands, targeting most lung cell populations, including itself (Fig. 2A), Among the most broadly acting ligands were *Agt*, *App*, and *Apoe* (Fig. 2C). Functional enrichment analysis (FEA) revealed, that the expression of the L-MSC-produced ligands was associated with pathways related to angiogenesis, cell migration, adhesion and chemotaxis, and ECM organization (Supplementary fig. 2B, Supplementary table 7).

**Figure 2.**
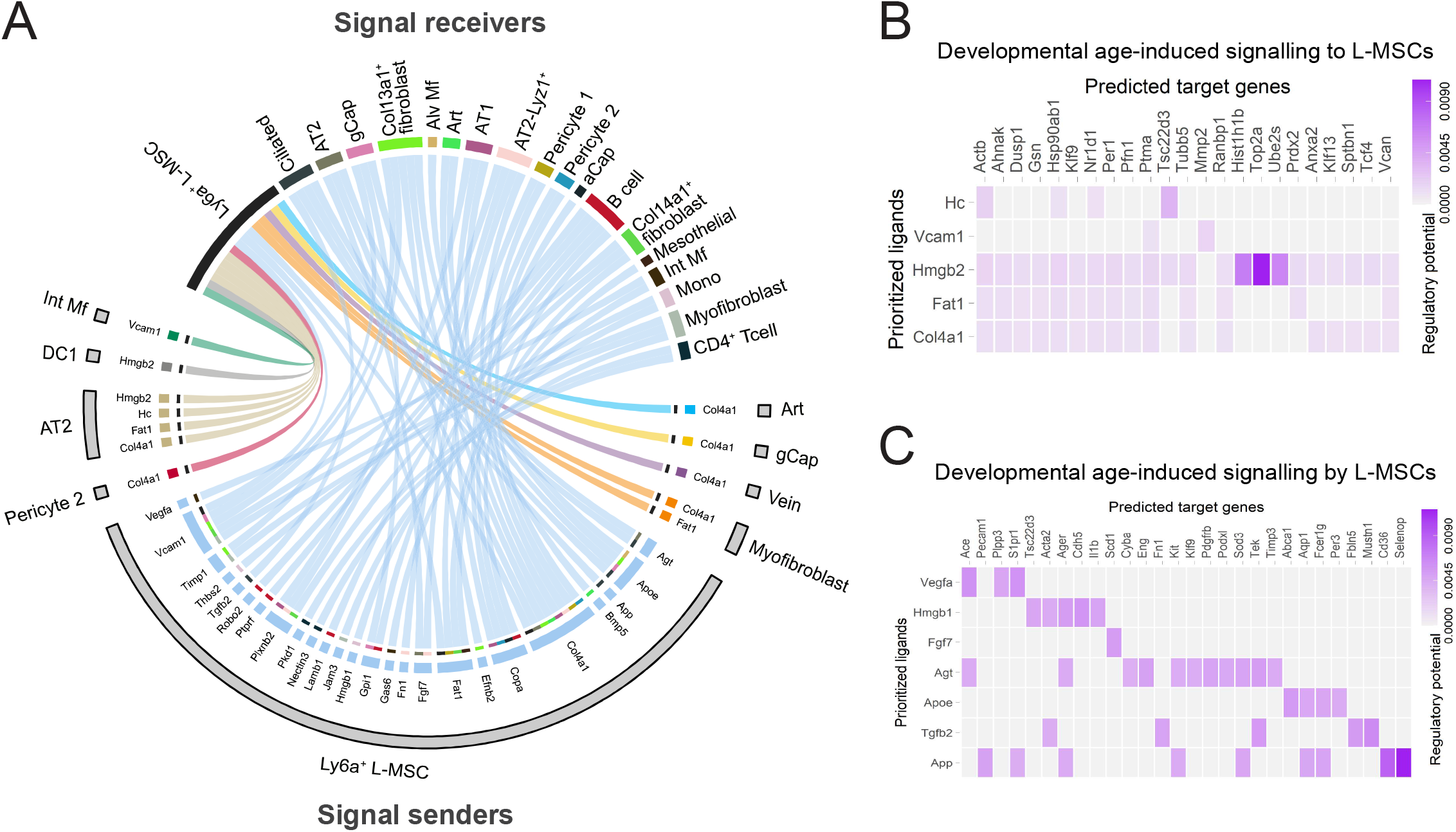
Age-associated gene expression and signalling in the developing *Ly6a*^+^ L-MSCs. **(A)** Circos plot showing inferred cell communications between *Ly6a*^+^ L-MSCs and other populations in the developing mouse lung. Cell communications associated with increasing developmental age are depicted. Cell types in the top right correspond to receiver populations with the largest expression changes in response to increasing age. These cell types are connected to the sender cell types expressing ligands predicted to promote this response. Ligands expressed by the same cell population are coloured the same. **(B)** Heatmap depicting predicted target genes for ligands most likely to be received by normally developing *Ly6a*^+^ L-MSC population as indicated in **(A)**. The intensity of expression is indicated as specified by the colour legend. **(C)** Heatmap depicting predicted target genes for ligands sent by *Ly6a*^+^ L-MSC population in normally developing lungs as indicated in **(A)**. The intensity of expression is indicated as specified by the colour legend.

### 3.3 The transcription profile and signaling activity of *Ly6a*^+^ L-MSC change significantly during postnatal lung development in response to hyperoxia

Hyperoxia induced an increase in proportion of *Ly6a*^+^ stromal cells as determined by scRNA-seq analysis at P14 (Fig. 1F, Supplementary table 3). This was consistent with increased proportion of LY6A^+^ stromal cells in hyperoxia-exposed lungs at P7 as measured by flow cytometry (Supplementary fig. 2C-D). In order to identify hyperoxia-induced changes in gene expression specific to *Ly6a*^+^ L-MSCs, we performed a DSA for both, *Ly6a*^+^ L-MSC population and non-progenitor *Ly6a*^−^stromal cells (Supplementary table 8). Hyperoxia-induced expression changes most distinctive of *Ly6a*^+^ L-MSCs are illustrated in Fig. 3A. Exposure to hyperoxia was associated with *Ly6a*^+^ L-MSCs - specific increase in expression of multiple pro-inflammatory (*Cxcl1*, *Ccl2*), as well as pro-fibrotic and anti-angiogenic (*Timp1*, *Serpina3n*) genes (Fig. 3A, Supplementary table 8). GSEA of hyperoxia-induced changes in gene expression revealed an activation in inflammatory pathways, as well as decrease in pathways associated with arterial development and morphogenesis (Fig. 3B, Supplementary table 9). When inspecting pathways altered by hyperoxia exclusively in *Ly6a*^+^, but not *Ly6a*^−^ stromal cells, activation of cytokine and chemokine signaling, cell cycle regulation, and senescence were most noticeable (Supplementary fig. 2E, Supplementary table 9).

**Figure 3.**
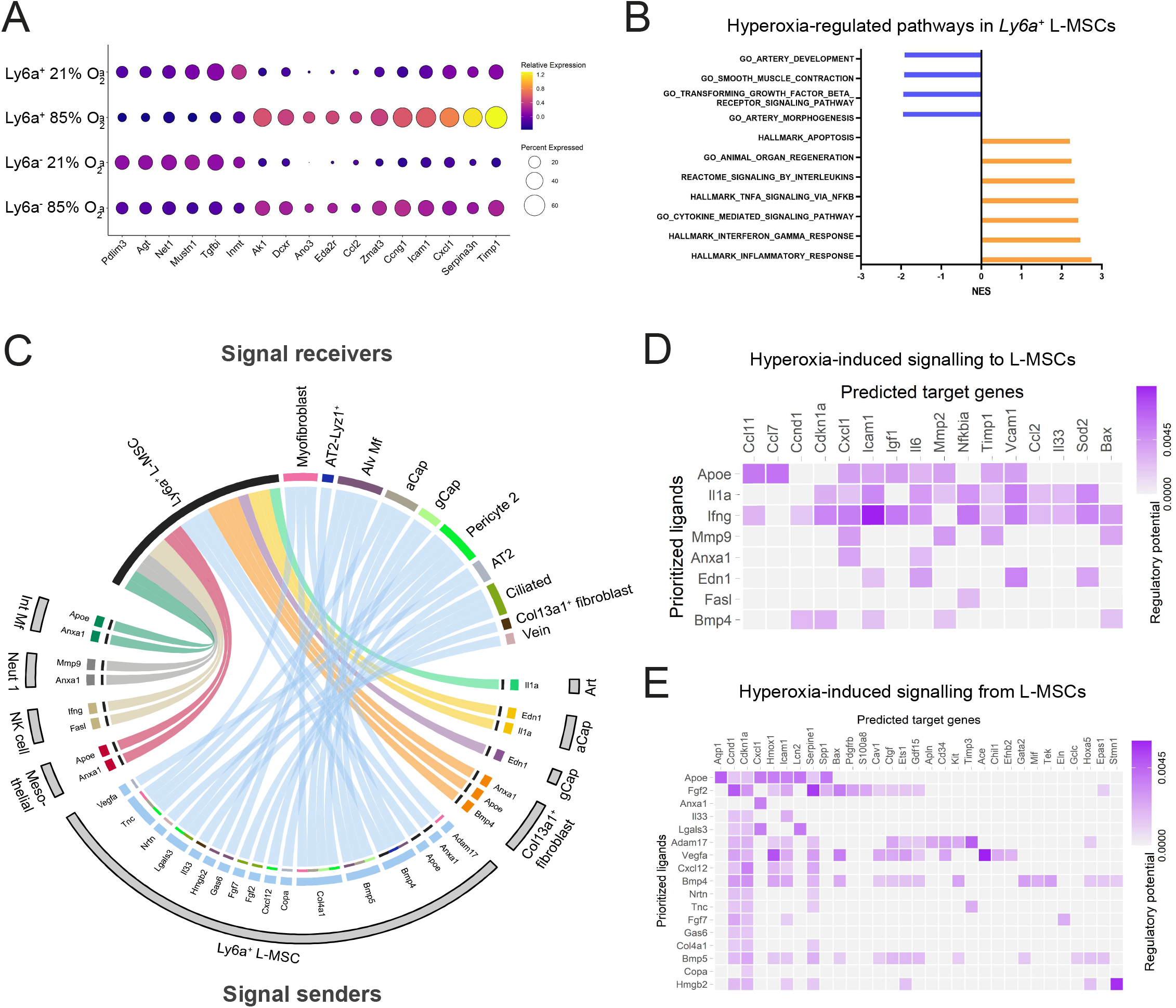
Hyperoxia-induced gene expression and signalling in the developing *Ly6a*^+^ L-MSCs. **(A)** Dotplot depicting the expression of markers specifically altered by hyperoxia exposure in *Ly6a*^+^ and *Ly6a*^−^ cells in the developing mouse lung. **(B)** Selected hyperoxia-regulated signalling pathways in the *Ly6a*^+^ L-MSC cluster identified by gene set enrichment analysis (GSEA). All terms are significantly enriched (adjusted p-value < 0.05). Normalized enrichment scores (NES) values were computed by gene set enrichment analysis on fold change-ranked genes. **(C)** Circos plot showing inferred cell communications between *Ly6a*^+^ L-MSCs and other populations in the developing mouse lung. Cell communications induced by exposure to hyperoxia are depicted. Cell types in the top right correspond to receiver populations with the largest expression changes in response to hyperoxia. These cell types are connected to the sender cell types expressing ligands predicted to promote this response. Ligands expressed by the same cell population are coloured the same. **(D)** Heatmap depicting predicted target genes for ligands most likely to be received by *Ly6a*^+^ L-MSC population in hyperoxia as indicated in **(C)**. The intensity of expression is indicated as specified by the colour legend. **(E)** Heatmap depicting predicted target genes for ligands sent by *Ly6a*^+^ L-MSC population in hyperoxia as indicated in **(C)**. The intensity of expression is indicated as specified by the colour legend.

To further understand the faith of *Ly6a*^+^ L-MSCs in hyperoxia-induced injury, we performed a cell communication analysis using the NicheNet tool, inferring hyperoxia-induced cellular communications[23,28] (Fig. 3C, Supplementary fig. 6-7, Supplementary table 10). *Ly6a*^+^ L-MSCs in hyperoxia-exposed lungs received signals from several cell populations, including immune cells, capillary and arterial endothelial cells, mesothelial cells and *Col13a1*^+^ fibroblasts (Fig. 3C). Further, we inferred genes in *Ly6a*^+^ L-MSCs most likely to be targeted by the received signals (Fig. 3D). Multiple ligands, such as *Apoe*, *Il1a*, *Ifng* and *Mmp9* were predicted to target the expression of pro-inflammatory, pro-fibrotic and anti-angiogenic genes discussed above, including *Timp1*, *Cxcl1* and *Icam1* (Fig. 3D). Expression of these target genes was elevated in *Ly6a*^+^ L-MSCs by hyperoxia exposure (Fig. 3A). Finally, ligands produced by *Ly6a*^+^ L-MSCs affected multiple cell populations, including alveolar macrophages, ciliated and AT2 cells, capillary and vein endothelium and other stromal populations. Among the most broadly acting ligands produced by *Ly6a*^+^ L-MSCs were *Bmp4*, *Bmp5*, *Col4a1* and *Tnc* (Fig. 3A). Inferred target genes in receiving cells targeted by majority of these ligands included *Ccnd1*, *Cdkn1a*, *Icam1* and *Hmox1* (Fig. 3E). According to FEA, expression of the L-MSC-produced ligands were associated with pathways related to vessel morphogenesis, epithelial cell proliferation, cell chemotaxis, and immune homeostasis and response (Supplementary fig. 2F, Supplementary table 11).

### 3.4 Murine LY6A^+^ L-MSCs localize to perivascular regions of the developing lung

Next, we aimed to localize the *Ly6a*^+^ L-MSCs in the developing lung using FISH. L-MSCs were identified as *Ly6a*^+^/*Col14a1*+ cells. L-MSCs in both, normally and aberrantly-developing lungs localized to perivascular regions of large vessels with more double-positive cells observed in hyperoxia-exposed lungs (Fig. 4A).

**Figure 4.**
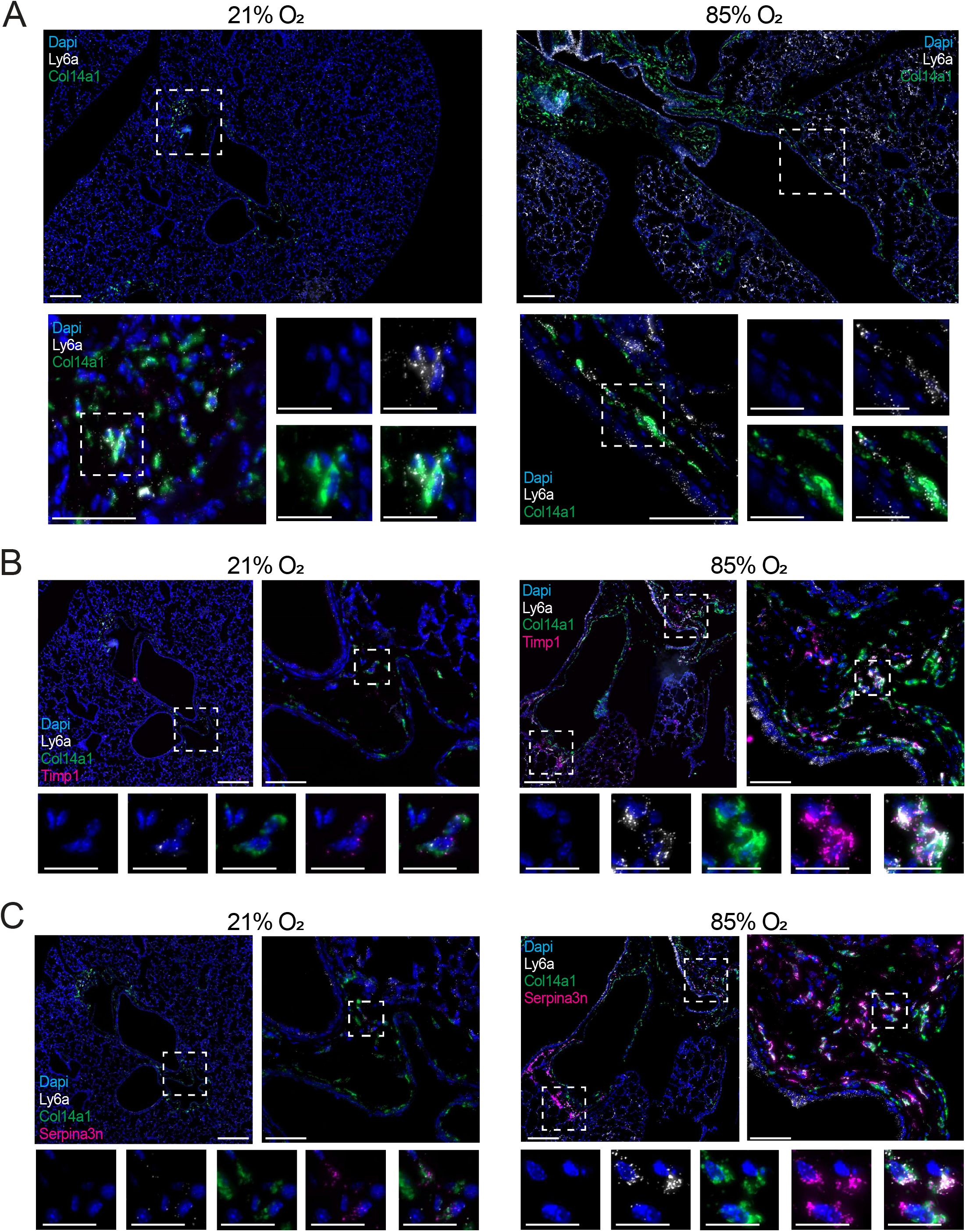
Identification of *Ly6a*^+^ L-MSCs in the developing lung. **(A)** Fluorescent RNA *in situ* hybridization showing localization of L-MSCs identified by the co-expression of *Ly6a* (white) and *Col14a1* (green) in lungs of room air (21% O_2_) or hyperoxia (85%O_2_)-exposed developing mice. Scale bar = 200μm for low-magnification (5×, top panels) windows, 50μm for higher-magnification (40×, bottom left panels) windows, and 20μm for high-magnification (63×, bottom right panels) windows. Four 14-days old animals/group were analysed. Expression levels in Dotplot are presented as log(TP10k+1) values. Log(TP10k+1) corresponds to log-transformed UMIs per 10k. **(B)** Fluorescent RNA *in situ* hybridization showing co-expression of *Ly6a* (white), *Col14a1* (green), and *Timp1* (pink) in lungs of room air (21% O_2_) or hyperoxia (85%O_2_)-exposed developing mice. Scale bar = 200μm for low-magnification (5×, top left) windows, 50μm for higher-magnification (40×, top right) windows, and 20μm for high-magnification (63×, bottom panels) windows. Four 14-days old animals/group were analysed. **(C)** Fluorescent RNA *in situ* hybridization showing co-expression of *Ly6a* (white), *Col14a1* (green), and *Serpina3n* (pink) in lungs of room air (21% O_2_) or hyperoxia (85%O_2_)-exposed developing mice. Scale bar = 200μm for low-magnification (5×, top left) windows, 50μm for higher-magnification (40×, top right) windows, and 20μm for high-magnification (63×, bottom panels) windows. Four 14-days old animals/group were analysed.

Additionally, we aimed to validate some of the novel normoxic and hyperoxic L-MSC markers as suggested by scRNA-seq analysis (Fig. 1C, Fig. 3A). *Ly6a*^+^ L-MSCs were co-stained for the hyperoxia-associated markers *Timp1* and *Serpina3n* (Fig. 4B and 4C, respectively). In both instances triple-positive cells were observed in the regions adjacent to large vessels (highlighted by white squares in low-magnification panels). These cells were not only more abundant in the lungs from BPD mice, but the expression levels of both, *Timp1* and *Seprina3n* were increased in the diseased lungs (see higher-magnification panels Fig. 4B-C).

### 3.5 Hyperoxia exposure does not impact clonal or differentiation potential of LY6A^+^ L-MSCs

In order to verify their progenitor cell-like properties, we isolated and studied LY6A^+^ L-MSCs from healthy and hyperoxia-exposed developing mouse pups. An arrest in lung development was induced by exposing newborn mouse pups to normobaric hyperoxia (85% O_2_) (Fig. 5A). CD31-/CD45^−^/EpCAM^−^/LY6A^+^ L-MSCs were isolated from seven days-old healthy (21% O_2_-exposed) or diseased (85% O_2_-exposed) mouse pups (Fig. 5B) and examined for the hallmarks of the MSC phenotype *in vitro*. While lungs of hyperoxia-exposed pups consistently yielded higher numbers of LY6A^+^ L-MSCs (Fig. 5B), no differences in the appearance (Fig. 5C), differentiation capacity (Fig. 5C), expression of surface markers (Fig. 5D), or clonal abilities (Fig. 5E) were observed between the cells isolated from healthy and diseased animals. LY6A^+^ L-MSCs isolated from both healthy and hyperoxia-exposed mice had a fibroblast-like appearance and expressed classical markers of MSCs *in vitro* (Fig. 5C-D). In order to investigate their differentiation capacity, LY6A^+^ L-MSCs were induced to differentiate along the osteogenic, chondrogenic, and adipogenic lineages. Both normoxia and hyperoxia-derived LY6A^+^ L-MSCs produced osteogenic and chondrogenic matrix (Fig. 5C). However, only a single sample of normoxia-derived LY6A^+^ L-MSCs produced a small number of adipocytes, and no lipogenic differentiation was observed in hyperoxia-derived LY6A^+^ L-MSCs (data not shown). Postnatal hyperoxia exposure had no effect on colony-forming capacity of LY6A^+^ L-MSCs as assessed by single-cell plating colony-forming assay. Both normoxia and hyperoxia-derived LY6A^+^ L-MSCs produced colonies of various sizes. While larger colonies consisted of fibroblast-like spindle-shaped cells, smaller colonies were formed by cells with a large cytoplasm (Fig. 5E). Inconsistent differentiation capacity and colony formation might suggest a heterogeneous nature of the LY6A^+^ L-MSCs population.

**Figure 5.**
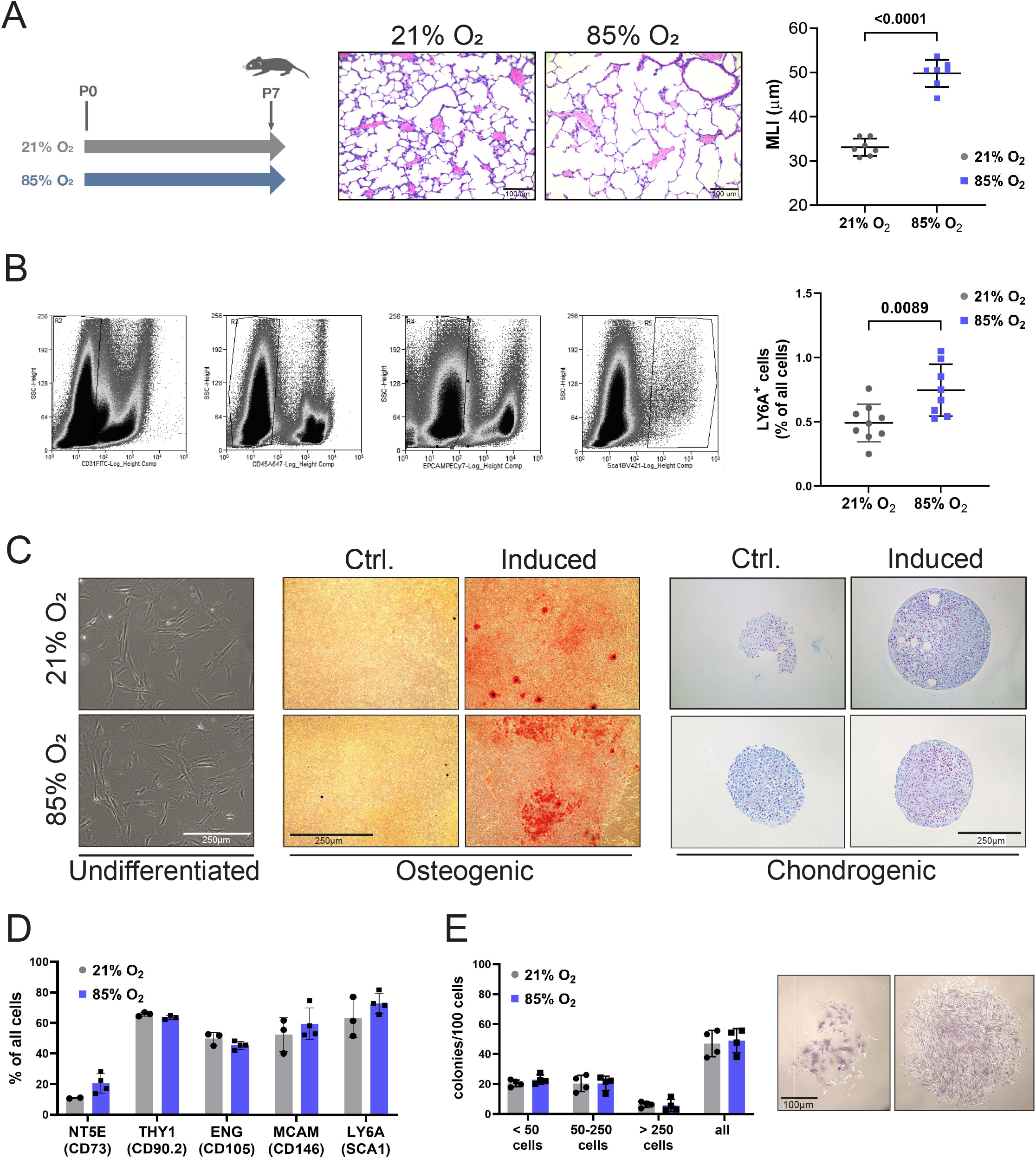
Characterization of LY6A^+^ L-MSCs in normal and impaired mouse lung development. **(A)** Mouse pups were exposed to room air (21% O_2_, grey) or hyperoxia (85%O_2_, blue) from P1 onwards. Mice were harvested on postnatal day (P)7. Representative histological sections from lungs developing in 21% O_2_ or 85% O_2_. Lung morphometry was quantified by the mean linear intercept (MLI) measurement. n = 7 animals/group. Scale bar = 100μm. **(B)** LY6A^+^ L-MSCs were identified by flow cytometry as CD45-AF647^−^/CD31-FITC^−^/CD326(EPCAM)-PeCy7^−^/LY6A(SCA1)-BV421^+^ cells and their proportion in lung homogenates was quantified. n = 8-9 animals/group. **(C)** Representative images of undifferentiated LY6A^+^ L-MSCs and LY6A^+^ L-MSCs differentiated towards osteogenic and chondrogenic lineages and stained with Alizarin Red S or Alcian Blue, respectively. Scale bar = 250μm. Experiments were performed in quadruplets. **(D)** Expression of routine MSCs surface markers in cultured LY6A^+^ L-MSCs isolated from room air (21% O_2_, grey bars) or hyperoxia-exposed (85%O_2_, purple bars) developing pups as determined by flow cytometry. n = 3-4 animals/group. **(E)** Quantification and representative images of colony formation of cultured LY6A^+^ L-MSCs isolated from room air (21% O_2_, grey bars) or hyperoxia-exposed (85%O_2_, purple bars). n = 4 animals/group. Scale bar =100μm. All data are presented as means ± SD. Statistical analyses were performed with GraphPad Prism 8.0. The presence of potential statistical outliers was determined by Grubbs’ test. Significance was evaluated by unpaired Student’s *t-*test for analysis in panels (A) and (B), and by multiple unpaired Student’s *t*-test with Holm-Šidák correction in panels (D) and (E). *P* values < 0.05 were considered significant and are depicted.

### 3.6 Cell culture alters the gene expression profile of LY6A^+^ L-MSCs

For therapeutic applications, MSCs are typically culture expanded, then frozen, over the short-, or long-term and thawed prior to administration. These various steps may alter the properties of the cell product. In order to understand changes in the L-MSCs expression profile induced by storage and culture, we performed a scRNA-seq analysis of cultured LY6A^+^ and LY6A-lung stromal cells isolated from seven days-old healthy (21% O_2_-exposed) or diseased (85% O_2_-exposed) mouse pups (Fig. 6A, Supplementary fig. 2C-D). We sequenced over 15,000 cultured CD31-/CD45-/EpCAM-/LY6A- and CD31-/CD45-/EpCAM-/LY6A^+^ cells and identified four distinct clusters (Fig. 6A-C, Supplementary tables 12-13). While normoxia and hyperoxia-derived LY6A-stromal cells contributed to all four clusters, very few LY6A^+^ cells could be found in clusters 2 and 3 (Fig. 6B). The presence of distinct clusters within the L-MSCs population is consistent with the heterogeneous phenotype of cultured L-MSCs described above (Fig. 5E). In line with this finding, the highest levels of routine MSC markers, such as *Thy1*, *Eng*, *Alcam* or *Mcam*, were found in the largest cluster 0, while very little expression was seen in the two smallest clusters (Fig. 6D). While still expressing routine MSC markers to some level, cluster 1 was characterized by its distinct expression of *Cck*, previously found to attenuate *p53*-mediated apoptosis in lung cancer [31] (Figure 3A-C). Cluster 2 was distinguished by the expression of pro-adipogenic markers, such as *Igfbp2* and *Col4a1*, as well as markers of myofibroblasts (Des) and alveolar epithelium (*Krt8* and *Prnp2*) (Fig. 6C, Supplementary table 13). Cluster 3 was characterized by the expression of multiple osteogenic markers, including *Cryab*, *Postn* and *Ngfr*. Interestingly, the expression of both *Postn*, as well as another cluster 3 marker *Col18a1*, was previously reported in BPD patients and hyperoxia-exposed developing mice [32,33].

**Figure 6.**
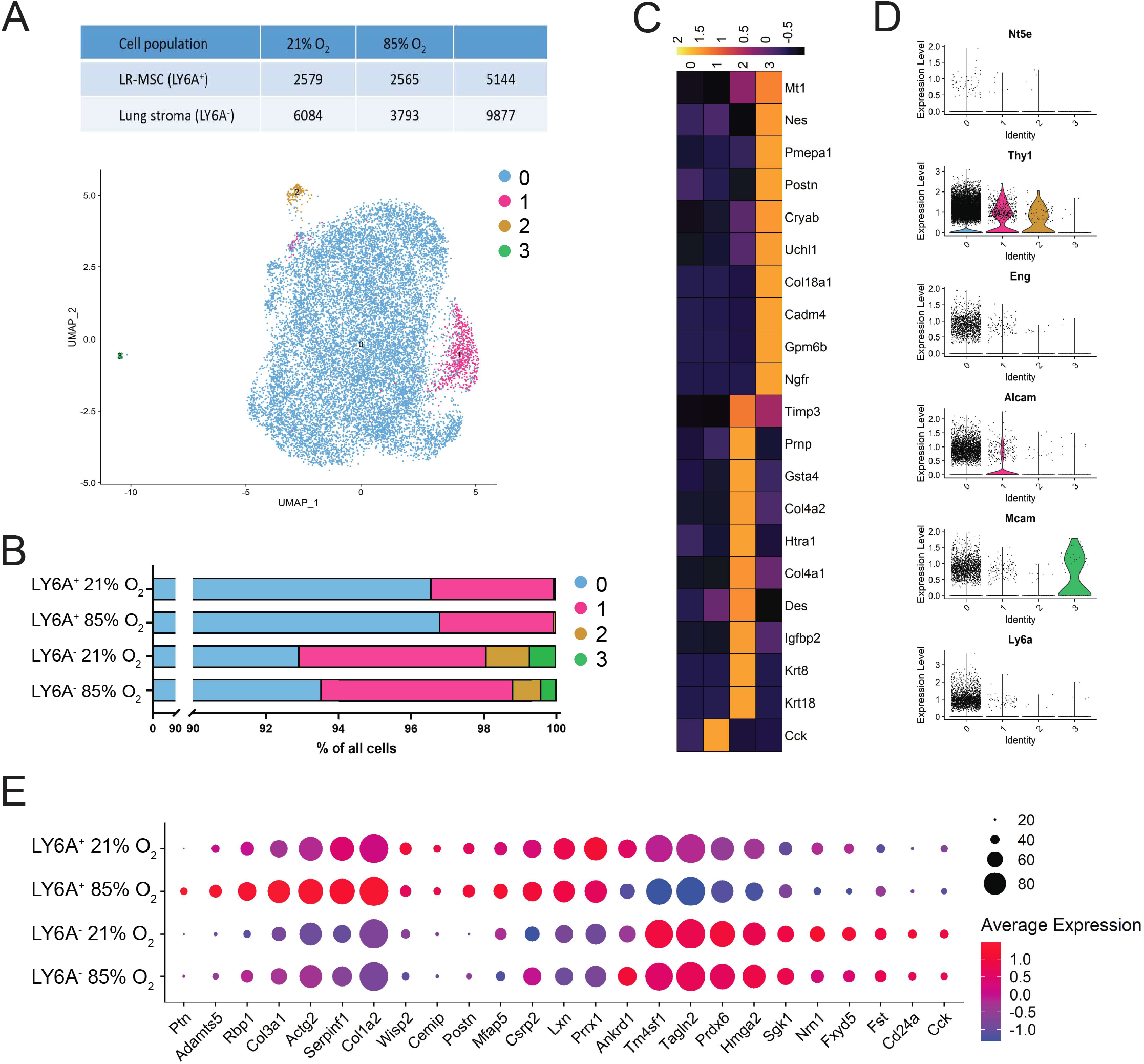
Gene expression profile of cultured normoxia and hyperoxia-derived LY6A^+^ L-MSCs. **(A)** LY6A^+^ and LY6A^−^ stromal cells isolated from lungs of room air (21% O_2_) or hyperoxia (85%O_2_)-exposed developing mice were frozen, cultured and sequenced at passage 3. n = 3 animals/group. scRNA-seq identified four clusters of cultured LY6A^+^ and LY6A^−^ stromal cells. **(B)** Relative distribution of room air (21% O_2_) or hyperoxia (85%O_2_)-derived LY6A^+^ and LY6A^−^ cells to the four different clusters. n = 3 animals/group. **(C)** Heatmap of top ten most differentially expressed genes across clusters depicted in panel (A). **(D)** Violin plots depicting expression of routine MSC markers in cultured stromal populations. **(E)** Dotplot depicting expression of oxygen-specific markers in LY6A^+^ and LY6A^−^ cultured lung stromal cells. Expression values in Heatmap and violin plots represent Z-score-transformed log(TP10k+1) values. Expression levels in Dotplot and UMAP plot are presented as log(TP10k+1) values. Log(TP10k+1) corresponds to log-transformed UMIs per 10k.

Next, we aimed to identify the best markers for cultured L-MSCs (Supplementary tables 14-18). We compared the gene expression profiles of LY6A^+^ and LY6A^−^ stromal cells (Supplementary tables 15-18) and identified differentially expressed genes between normoxia- and hyperoxia-derived subsets of these populations (Supplementary tables 17-18). In comparison to LY6A^−^ cells, LY6A^+^ cells were characterized by high expression of *Actg2, Col1a2*, *Serpinf1*, *Prrx1* and *Lxn*, and by low expression of smooth muscle cell (SMC) marker *Tagln2*[34], alveolar progenitor marker *Tm4sf1*[35], and *Prdx6* (Fig. 6E, Supplementary tables 15-16). From these markers hyperoxia exposure further specifically increased the expression of *Actg2*, and decreased the expression of *Tagln2*, *Tm4sf1* and *Prdx6* in LY6A^+^ cells. Hyperoxic LY6A^+^ cells were additionally distinguished by expression of *Ptn*, *Adamts5*, *Rbp*, and *Col3a1* (Fig. 6E, Supplementary table 17). Expression of *Prrx1* and *Serpinf1* is known to favour an osteogenic phenotype, and *Serpinf1* is known to inhibit adipogenesis[36,37].

In order to identify L-MSCs expression patterns maintained after cell culture and storage, we next compared expression of the most promising markers of *Ly6a*^+^ L-MSCs in both, *in situ* and *in vitro* datasets from cells isolated at P7 (Supplementary fig. 2G-H). This analysis revealed that a large portion of the expression profile characteristic for *Ly6a*^+^ L-MSCs *in situ* (Supplementary fig. 2G-H) is lost when cells are frozen and cultured, including the expression of promising markers, such as *Lum*, *Ptn*, *Dcn*, or *Pi16* (Supplementary fig. 2H). Furthermore, while the expression pattern of some markers, such as *Serpina3n* or *C3* persisted in cultured cells, the portion of the cells expressing the gene was diminished (Fig. 7A-B). The most suitable *in situ* or *in vitro*-specific identifying markers of *Ly6a*^+^ L-MSCs are depicted in Fig. 4A-B. Among the most stable markers of *Ly6a*^+^ L-MSCs, resistant to changes induced by culture, were *Serpinf1* and *Postn* (Fig. 7A-B, Supplementary fig. 2G-H). In order to confirm the viability of *Serpinf1* as potential novel marker for L-MSCs we performed FISH in developing lungs at P14. Triple-positive cells could be found in lungs of both, normoxic and hyperoxic mice (Fig. 7C). No differences were apparent in *Serpinf1* expression intensity between the two groups.

**Figure 7.**
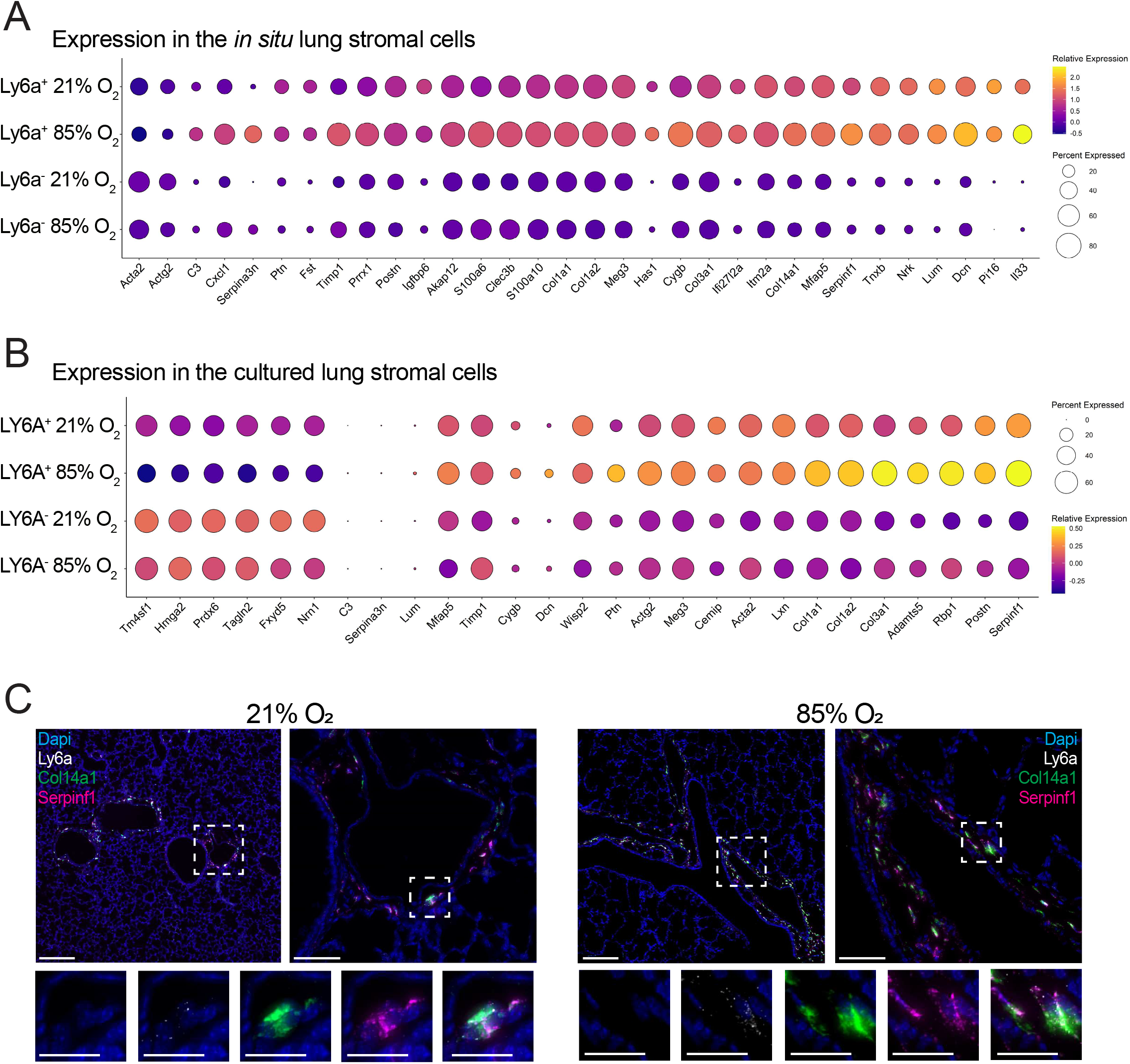
Identification of novel markers for *in situ* and cultured *Ly6a*^+^ L-MSCs. **(A)** Identifying markers were first established in the *in situ*, or cultured *Ly6a*^+^ and *LY6A*^−^ lung stromal cells based on Supplementary figures 1G-H. Dotplot depicts the expression levels of those markers, most suitable for identification of *LY6A*^+^ and *LY6A*^−^ lung stromal cells *in situ* in normoxic or hyperoxic animals at P7. (**B**) Identifying markers were first established in the *in situ*, or cultured *LY6A*^+^ and *LY6A*^−^ lung stromal cells based on Supplementary figures 1G-H. Dotplot depicts the expression levels of those markers, most suitable for identification of normoxia-derived and hyperoxia-derived LY6A^+^ and LY6A^−^ lung stromal cells in culture. **(C)** Fluorescent RNA *in situ* hybridization showing co-expression of *LY6A* (white), *Col14a1* (green), and *Serpinf1* (pink) in lungs of room air (21% O_2_) or hyperoxia (85%O_2_)-exposed developing mice. Scale bar = 200μm for low-magnification (5×, top left) windows, 50μm for higher-magnification (40×, top right) windows, and 20μm for high-magnification (63×, bottom panels) windows. Four 14-days old animals/group were analysed.

Finally, new expression patterns arose particularly in hyperoxia-derived *Ly6a*^+^ L-MSCs after cell culture. While a high *Ptn*, *Lum*, *Dcn, Col3a2* and *Col14a1* expression was initially characteristic of both, hyperoxia and normoxia-derived *Ly6a*^+^ L-MSCs, in cultured L-MSCs this was true only for the hyperoxia-derived *Ly6a*^+^ L-MSCs (Supplementary fig. 2G-H). This expression pattern denotes, that not only does the L-MSC transcriptome change in culture, but that the cells isolated from lungs of diseased mice tend to retain their expression profile and, potentially, progenitor-like nature longer.

## DISCUSSION

Our current knowledge regarding the identity and properties of tissue resident MSCs remains limited. Most studies analyse L-MSCs in culture after isolation with one, or several MSC markers. However, no explicit rules regarding which markers represent the L-MSC population the best exist to date. The progenitor-like characteristics of these cells have been established in culture [13,14], but it is not yet known why L-MSCs fail to prevent the lung injury or restore damage in the lung. While L-MSCs were previously found in bronchoalveolar lavage of BPD patients [50], it is not known whether this is due to increased apoptosis and subsequent shedding from the lung or is a sign of activation and proliferation of L-MSC and hence of increased numbers in BPD patients. Here, we provide an extensive scRNA-seq based analysis of L-MSCs in developing mouse lung, as well as in culture. We characterize the changes in trancriptomic profile induced in L-MSCs by developmental age, exposure to hyperoxia, and culture. Our study further provides an insight into communication between L-MSCs and other cell populations in the normally and abnormally developing lung. Finally, we propose novel markers for identification of L-MSCs in the developing lung.

The use of omics approaches to study tissue-specific MSCs *in vivo* has been previously proposed [38]. In the study presented here, we utilize scRNA-seq to study L-MSCs immediately after isolation (*in situ*) without confounding procedures, such as FACS, cell culture and storage, and hence preserve the in vivo activation status of the different lung populations as much as possible. We selected *Ly6a* to identify L-MSCs for 2 reasons: i) *Ly6a* is one of the most commonly used L-MSC markers and its expression has been shown in specific progenitor-like populations, ii) *Ly6a* was the only known MSC marker forming a visible subcluster within the lung mesenchyme of early postnatal mouse pups. We identified novel markers of L-MSCs, including *Lum*, *Serpinf1*, and *Dcn*. Next, we showed how the L-MSC’s transcriptome changes during the course of normal lung development and in hyperoxia, and explored the communication between L-MSCs and other lung cell populations.

Hyperoxia-exposure, used as a model for BPD, was associated in *LY6A*^+^ L-MSCs with increased expression of multiple pro-inflammatory (*Cxcl1*, *Ccl2*), pro-fibrotic and anti-angiogenic (*Timp1*, *Serpina3n*) genes. Similarly, increased expression of both, *Timp1* and *Ccl2* was previously reported in hyperoxia-exposed rodents [39,40], and in plasma [41] or tracheal aspirates (TA) [42] of BPD patients. *Timp1* expression was further increased in fibrotic foci in chronic BPD [43] and in the lungs of ventilated newborns [44]. GSEA further confirmed the activation of inflammatory and pro-fibrotic pathways, and a decrease in sprouting angiogenesis and vessel morphogenesis in the hyperoxia-exposed developing lungs. To further explore the role L-MSCs play in cell signaling during the development, we performed a cell communication inference analysis. L-MSCs in healthy developing lungs received ligands secreted mainly from endothelial, immune and other stromal cells. L-MSCs signalled back to the majority of lung cell populations with a selected set of ligands (Fig. 2). Upon hyperoxia exposure, L-MSCs received ligands primarily from immune and endothelial cells, including *Il1a*, *Mmp9*, *Ifng*, and *Fasl* (Fig. 3). Interestingly, multiple ligands received by L-MSCs were predicted to target the expression of pro-inflammatory, pro-fibrotic and anti-angiogenic genes increased in hyperoxia-exposed L-MSCs, such as *Timp1*, *Cxcl1* and *Icam1*. IFNγ and MMP9, which target the expression of both *Timp1* and *Cxcl1*, were previously implicated in development of alveolar hypoplasia [45] and an increased expression of IFNγ was reported in TA of BPD patients [27,46]. Development of BPD was also associated with increased TA and plasma protein levels of ICAM1 [47,48]. IL1A was also shown to induce an inflammatory phenotype in lung fibroblasts [49]. Additionally, *Fasl*^+^ immune cells were shown to induce fibroblast cell death [50,51], and its overexpression was associated with alveolar apoptosis and disturbed alveolar and vascular development [52].

Next, we investigated how the L-MSCs’ transcriptome changed due to culture and storage, both necessary steps for the preparation of a cell therapeutic product. ScRNA-seq analysis revealed, that following isolation, storage and culture, most L-MSCs retain the expression of MSC markers, including *Ly6a*^+^. Cultured L-MSCs showed moderate ability to differentiate into chondrocytes and osteoblasts. However, we observed only one instance of successful differentiation along the adipogenic lineage, consistent with previous studies of L-MSCs in developing rats [13]. Inconsistent differentiation capacity could be attributed to heterogeneity within the L-MSC population as indicated by the variable size and morphology of L-MSC-derived colonies (Fig. 5). Importantly, such heterogeneity could indicate the existence of L-MSCs with varying progenitor-like capabilities, most likely impacting their therapeutic efficacy. Further, more detailed characterization of different L-MSCs subpopulations might be necessary in order to prepare a superior therapeutic product. ScRNA-seq revealed considerable changes in the transcriptome of L-MSCs in culture, implying that the cells studied and maintained *in vitro* for the purposes of therapeutic interventions are appreciably altered compared to L-MSCs *in situ* (Fig. 6). Interestingly, we observed that the culture-induced transcription changes are less pronounced in L-MSCs derived from hyperoxia-exposed animals. This might suggest that hyperoxia primes L-MSCs to maintain certain characteristics, potentially in an attempt to trigger a repair mechanism. While the organism’s own resident L-MSCs fail to prevent the hyperoxia-induced lung damage, a therapeutic use of injury-primed L-MSCs might be more beneficial than L-MSCs from healthy individuals. Interestingly, tissue origin and microenvironment were shown to significantly impact the behaviour and therapeutic efficacy of MSCs [53,54]. Moreover, conditioned media from BM-MSCs exposed *ex vivo* to hyperoxia exhibited superior therapeutic effects in the hyperoxia-induced rat BPD model when compared to media from BM-MSCs which were not pre-conditioned [55].

The localization of L-MSCs in the developing lungs has not yet been described. Here, we localized the L-MSC cells in the perivascular regions of both, heathy and diseased developing lungs by FISH. The *Ly6a*^+^ L-MSCs in the hyperoxia-exposed lungs co-expressed *Timp1* and *Serpina3e*, confirming the results of scRNA-seq analysis. Finally, as *Ly6a* is not expressed in human tissues, we aimed to identify additional markers to label L-MSCs, both *in situ* and *in vitro*. *Lum*, identified as marker of L-MSCs *in situ*, is known to be produced by MSCs. Within the lung, it’s expression was localized to peripheral lung and vessel walls [56]. While scRNA-seq revealed *Lum* as a promising L-MSCs marker *in situ*, it’s expression in culture was preserved only in a small fraction of L-MSCs isolated from hyperoxia-exposed animals (Fig. 7A). In comparison, the expression of *Serpinf1* was well preserved *in vitro*, with the expression slightly increased in hyperoxic cells. Interestingly, *Serpinf1* expression was previously reported to be increased in hyperoxia-exposed newborn mice and *Serpinf1*^−/−^ animals were protected from hyperoxia-induced lung injury [57]. *Serpinf1* is also known as an anti-angiogenic and anti-migratory marker associated with aging MSCs [57,58]. *In situ*, *Serpinf1* colocalized well with *Ly6a*^+^/*Col14a1*^+^ cells in both healthy and diseased lungs, suggesting *Serpinf1* as promising new marker for L-MSCs (Fig. 7).

To our knowledge, this is the first detailed report studying the characteristics and behaviour of L-MSC *in situ* and *in vitro*, during both health and disease. We unravelled the transcriptome and cellular communication of this lung resident cell population by scRNA-seq in order to mechanistically understand its endogenous repair capabilities, as well as its potential use as an exogenous cell therapeutic product. In addition, we have established several markers that can be used to identify L-MSC *in vitro* and *in vivo*, both in healthy and diseased lungs. Additional studies will be needed to further unravel the heterogeneity of this population, as well as their therapeutic capabilities.

## ACKNOWLEDGMENTS

This study was supported by the Canadian Institutes of Health Research (CIHR), the German Research Foundation (Deutsche Forschungsgemeinschaft), the Frederick Banting and Charles Best Doctoral Scholarship, the Finnish Foundation for Pediatric Research, the Finnish Sigrid Juselius Foundation, the Canadian Lung Association - Breathing as One, and the Molly Towel Perinatal Research Foundation.

## DICLOSURE OF POTENTIAL CONFICT OF INTERESTS

The authors declare no conflicts of interest, financially or otherwise.

## AUTHOR CONTRIBUTIONS

IM: Conception and design, collection and assembly of data, data analysis and interpretation, manuscript writing, final approval of manuscript. DPC: Collection and assembly of data, data analysis and interpretation, manuscript writing, final approval of manuscript. CCD: Collection and assembly of data, data analysis and interpretation, manuscript writing, final approval of manuscript. FL: Collection and assembly of data, data analysis and interpretation, manuscript writing, final approval of manuscript. MH: Collection and assembly of data, final approval of manuscript. SMH: Collection and assembly of data, final approval of manuscript. SZ: Collection and assembly of data, final approval of manuscript. OC: Financial support, administrative support, final approval of manuscript. BCV: Financial support, administrative support, final approval of manuscript. BT: Conception and design, financial support, administrative support, final approval of manuscript.

## DATA AVAILABILITY STATEMENT

