## Supplemental Table 1 for "Single-cell RNA sequencing-based characterization of resident lung mesenchymal stromal cells in bronchopulmonary dysplasia"

**Supplementary table 1. Antibodies used for flow cytometry.**

| Antigen of interest | Conjugated fluorophore | Cat. No., company |
|---|---|---|
| CD16/32 (Fc block) | - | #553142, BD Biosciences, Mississauga, ON, Canada |
| CD31 | FITC | #558738, BD Biosciences, Mississauga, ON, Canada |
| CD45 | AF647 | #1660-31, Southern Biotech, Birmingham, AL, USA |
| CD326 (EpCAM) | Pe/Cy7 | #25-5791-80, ThermoFisher Scientific, Burlington, ON, Canada |
| LY-6A/E (SCA1) | BV421 | #744322, BD Biosciences, Mississauga, ON, Canada |
| CD31 | BV421 | #102424, BioLegend, SanDiego, CA, USA |
| CD73 | PE | #127206, BioLegend, SanDiego, CA, USA |
| CD105 | AF488 | #120406, Biolegend, SanDiego, CA, USA |
| CD34 | PE | #551387, BD Biosciences, Mississauga, ON, Canada |
| CD146 | AF488 | #562229, BD Biosciences, Mississauga, ON, Canada |
| CD90.2 | PB | #105324, BioLegend, SanDiego, CA, USA |
