## Supplementary Methods for "Single-cell RNA sequencing-based characterization of resident lung mesenchymal stromal cells in bronchopulmonary dysplasia"

### Method S1

##### Tissue digestion

Isolated mouse lungs were dissected into individual lobes, and digested at 37°C by gentleMACS™ Octo Dissociator in following enzyme mix: 2500U Collagenase I (Worthington Biochem., Lakewood, NJ, USA), 30U Neutral Protease (Worthington Biochem., Lakewood, NJ, USA), 500U Deoxyribonuclease (DNAse) I (Sigma-Aldrich, Oakville, ON, Canada) in 5ml of DPBS supplemented with $Mg^{2+}/Ca^{2+}$ (ThermoFisher Scientific, Burlington, ON, Canada). The following dissociation program was used: loop 6× (spin 300rpm, 10"; spin -300rpm, 10"); loop 2× (spin 150rpm, 5"; spin -150rpm, 5"); loop 2× (spin 20rpm, 5' 0"; spin -20rpm, 5' 0"); loop 6× (ramp 360rpm, 15"; ramp -360rpm, 15"). The resulting cell suspension was filtered through a 100 µm nylon mesh (ThermoFisher Scientific, Burlington, ON, Canada) and the enzymatic reaction was terminated by 0.9 mM EDTA (Invitrogen, Carlsbad, CA, USA).

### Method S2

##### Cell culture and storage

Following isolation and FACS, $CD31^-/CD45^-/EpCAM^-/LY6A^-$ and $CD31^-/CD45^-/EpCAM^-/LY6A^+$ lung cells from each mouse were plated separately in T25 culture flasks (ThermoFisher Scientific, Burlington, ON, Canada). Cells were cultured in 5ml growth media containing alpha-modified Minimum Essential Medium Eagle (αMEME) medium (Sigma-Aldrich, Oakville, ON, Canada), complemented with 20% FBS, 1% antibiotic-antimycotic (ThermoFisher Scientific, Burlington, ON, Canada), and 2mM L-Glutamine (ThermoFisher Scientific, Burlington, ON, Canada). Cell medium was exchanged 24 hours after isolation and in 48 hours intervals after that. After reaching 70% confluency, usually 4-5 days after isolation, cells were passaged to T75 culture flasks (Sarstedt, Nümbrecht, Germany). Cells were kept at 37°C with 5% $CO_2$ and 5% $O_2$. All cells were frozen in freezing media [20% FBS, 5% DMSO (Sigma-Aldrich, Oakville, ON, Canada), 6% (w/v) Hydroxyethylstarch (AK Scientific, Union City, CA, USA), 12.5% CPDA-1 in 0.56% NaCl] at passage 2 and stored in liquid nitrogen.

### Method S3

##### Colony formation assay

Passage 3 $CD31^-$/$CD45^-$/$EpCAM^-$/$LY6A^+$ L-MSCs were sorted using a BD LSR Fortessa (Beckton Dickinson Biosciences, Franklin Lakes, NJ, USA) at the Ottawa Hospital Research Institute (OHRI) StemCore facility to 96-well plates (Sarstedt, Nümbrecht, Germany) at a concentration of exactly one cell/well. Three technical replicates (three 96-well plates) were used for each sample. Cells were cultured in growth media for 11 days. Every 3-4 days, 50% of medium was gently refreshed. Cells were cultured in a cell culture incubator at 37°C with 5% $CO_2$ and 5% $O_2$. After 11 day of culture, cell colonies were fixed with ice-cold methanol and stained with Crystal Violet (Sigma-Aldrich, Oakville, ON, Canada). Colonies were photographed with EVOS XL Core microscope and counted according to their size.

#### Method S4

##### Osteogenic differnetiation

Cultured, passage 3 $CD31^-$/$CD45^-$/$EpCAM^-$/$LY6A^+$ L-MSCs were used for the osteogenic differentiation assay using the StemXVivo Osteogenic Supplement (Cat. No. #CCM009, R&D Systems, Minneapolis, MN, USA) according to the manufacturer's instructions. After 21 days of culture, cells were gently washed with dd$H_2O$, fixed with 4% PFA and stained with Alizarin Red S (Sigma-Aldrich, Oakville, ON, Canada) for 45 minutes in dark. Individual wells were photographed with EVOS XL Core (ThermoFisher Scientific, Burlington, ON, Canada) microscope.

#### Method S5

##### Adipogenic differnetiation

Cultured, passage 3 $CD31^-$/$CD45^-$/$EpCAM^-$/$LY6A^+$ L-MSCs were used for the adipogenic differentiation assay using the StemXVivo Adipogenic Supplement (Cat. No. #CCM011, R&D Systems, Minneapolis, MN, USA) according to the manufacturer's instructions. After 21 days of culture, cells were gently washed with dd$H_2O$, fixed with 10% buffered formalin (ThermoFisher

Scientific, Burlington, ON, Canada) and stained overnight in dark with Oil Red O (Sigma-Aldrich, Oakville, ON, Canada). Individual wells were photographed with EVOS XL Core microscope.

#### Method S6

##### Chondrogenic differnetiation

Cultured, passage 3 CD31$^-$/CD45$^-$/EpCAM$^-$/LY6A$^+$ L- were used for the chondrogenic differentiation assay using the StemXVivo Chondrogenic Supplement (Cat. No. #CCM006, R&D Systems, Minneapolis, MN, USA) according to the manufacturer's instructions. After 28 days medium was removed and cellular congregates were fixed with 10% buffered formalin. Cellular conglomerates were embedded in paraffin, sectioned at 4µm and stained with Alcian Blue. Tissue dehydration, paraffin embedding, sectioning and staining were performed by the University of Ottawa Louis Pelletier Histology Core Facility.

#### Method S7

##### Fluorescent in situ hybridization

RNA in situ hybridization was performed on 4% PFA-fixed paraffin embedded, 3µm tissue sections using RNAscope Multiplex Fluorescent Reagent Kit Version 2 (Advanced Cell Diagnostics, Newark, CA, USA) according to the manufacturer's instructions. Briefly, tissue sections were baked for 1 h at 60°C, deparaffinized and treated with hydrogen peroxide for 10 min at RT. Target retrieval was performed at 98°C for 15 min, followed by protease plus treatment at 40°C for 15 min. All probes were hybridized for 2 h at 40°C followed by signal amplification and developing of HRP channels. The following RNAscope probes were used in the study: 4-Plex Negative Control Probe-Mm (#321831), 4-Plex Positive Control Probe-Mm (#321811), Mm-Serpinf1-C4 (#310731-C4), Mm-Ly6a-C2 (#427571-C2), Mm-Serpina3n-C3 (#430191-C3), Mm-Timp1-C3 (#316841-C3), Mm-Col14a1 (#581941). All probes were optained from ACD Biotechne Minneapolis, MN, USA. TSA Plus fluorophores Fluorescein (1:100 dilution), Cyanine 3 (1:750 dilution), and Cyanine 5 (1:3000 dilution) (Perkin Elmer, Waltham, MA, USA) were used for signal detection. Sections were counterstained with DAPI and mounted with ProLong Gold Antifade Mountant (Invitrogen, Carlsbad, CA, USA). Tissue sections were scanned with

3DHISTECH Panoramic 250 FLASH II digital slide scanner at the Genome Biology Unit (Research Programs Unit, Faculty of Medicine, University of Helsinki, Biocenter Finland) at 40× magnification with extended focus and 7 focus levels.

#### Method S8

##### Multiplexing samples for scRNA-seq

Passage 3 $CD31^-/CD45^-/EpCAM^-/LY6A^-$ and $CD31^-/CD45^-/EpCAM^-/LY6A^+$ L-MSCc cells were plated in quadruplets in 96-well plates at concentration of 4.000 cells/well and reached approximately 80% confluency after 24 hours. Cells were lifted from the plates with 20µl of TrypLE-Express enzyme solution (ThermoFisher Scientific, Burlington, ON, Canada) containing 200nM anchor/200nM barcode solution (kindly provided by Prof. Zev Gartner from University of California, San Francisco). The lipid-modified DNA oligonucleotide (LMO) anchor and a unique "sample barcode" oligonucleotides were added to each sample in order to be multiplexed, with each sample receiving a different sample barcode. Samples were shaken gently and incubated in cell culture incubator at 37°C for 10 minutes. After 10 minutes, 150µl of TrypLE-Express/200nM co-anchor solution was added to each sample. Samples were shaken gently and incubated in cell culture incubator at 37°C for 5 minutes. After 5 minutes the reaction was neutralized by adding 50µl/sample of growth media and samples were transferred into a V-bottom 96-well plate (ThermoFisher Scientific, Burlington, ON, Canada) and pelleted at 500 × g for 5 minutes. Barcode-containing media was then removed, and the resulting cell pellet was washed twice with ice-cold 1% BSA (Sigma-Aldrich, Oakville, ON, Canada) in 1×PBS, and once in 1×PBS without BSA. After the last wash, pellets were resuspended in 1×PBS, counted and were pooled together at 1:1 ratio while maintaining the final concentration of 500-1000 cells/µl. Viability and cell counts were assessed using the EVE NanoEnTek automatic cell counter. Samples with viability ≥ 80% were further processed by 10× Chromium at the OHRI StemCore facility.
