## Supplementary figures and images for "Single-cell RNA sequencing-based characterization of resident lung mesenchymal stromal cells in bronchopulmonary dysplasia"

### Supplementary Figure 1

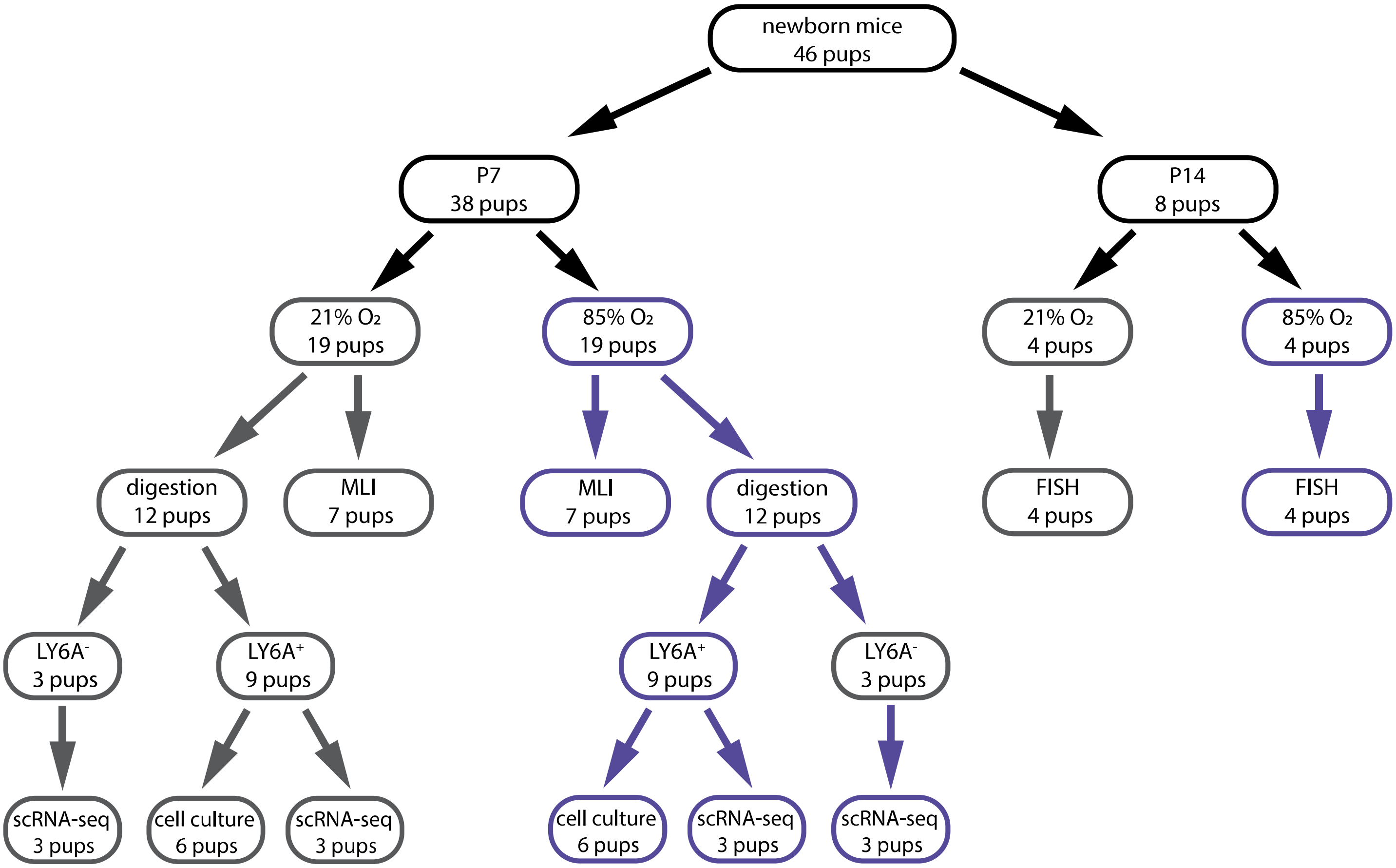


Supplementary Figure 1

### Supplementary Figure 2

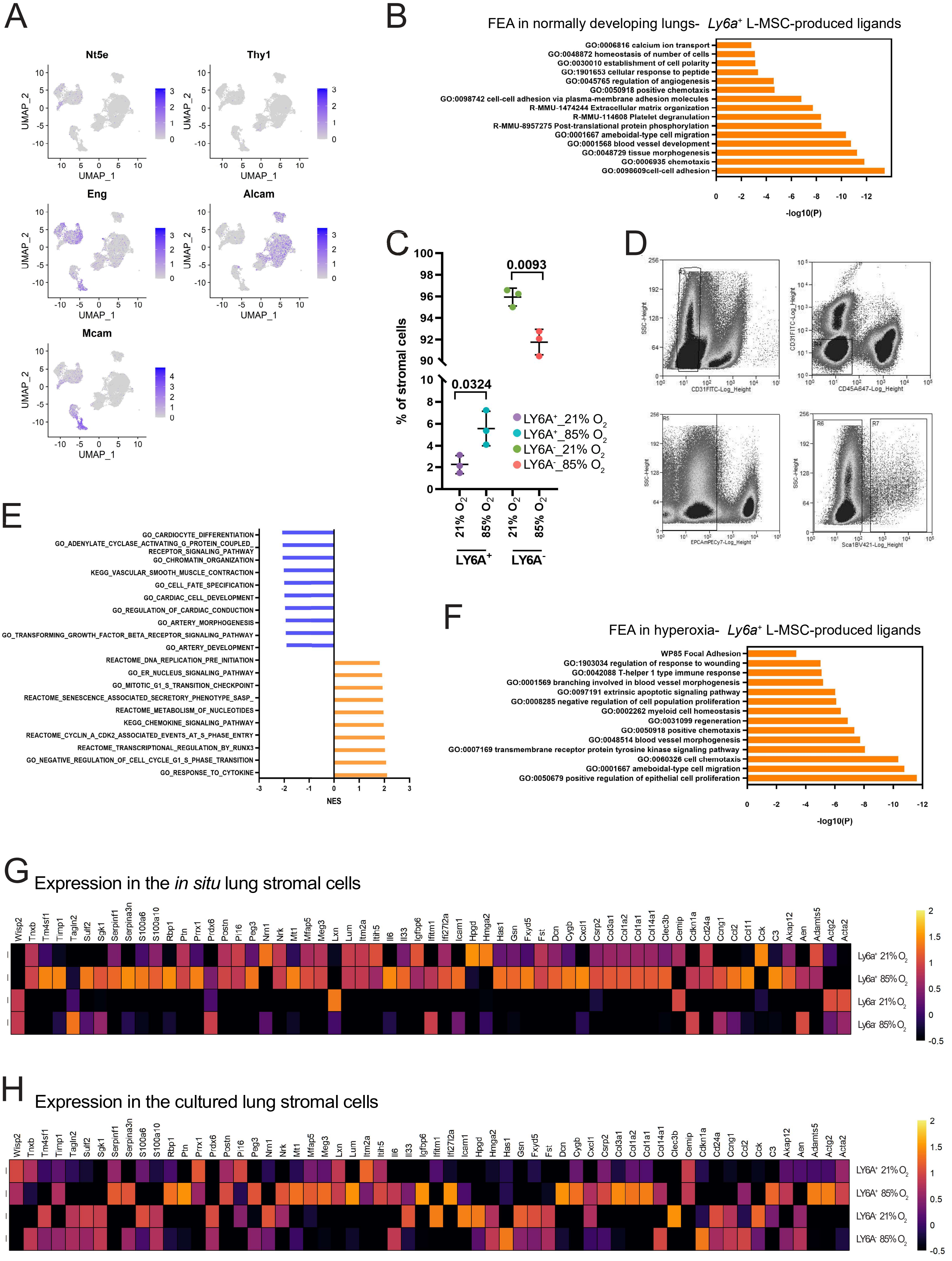


Supplementary Figure 2

### Supplementary Figure 3

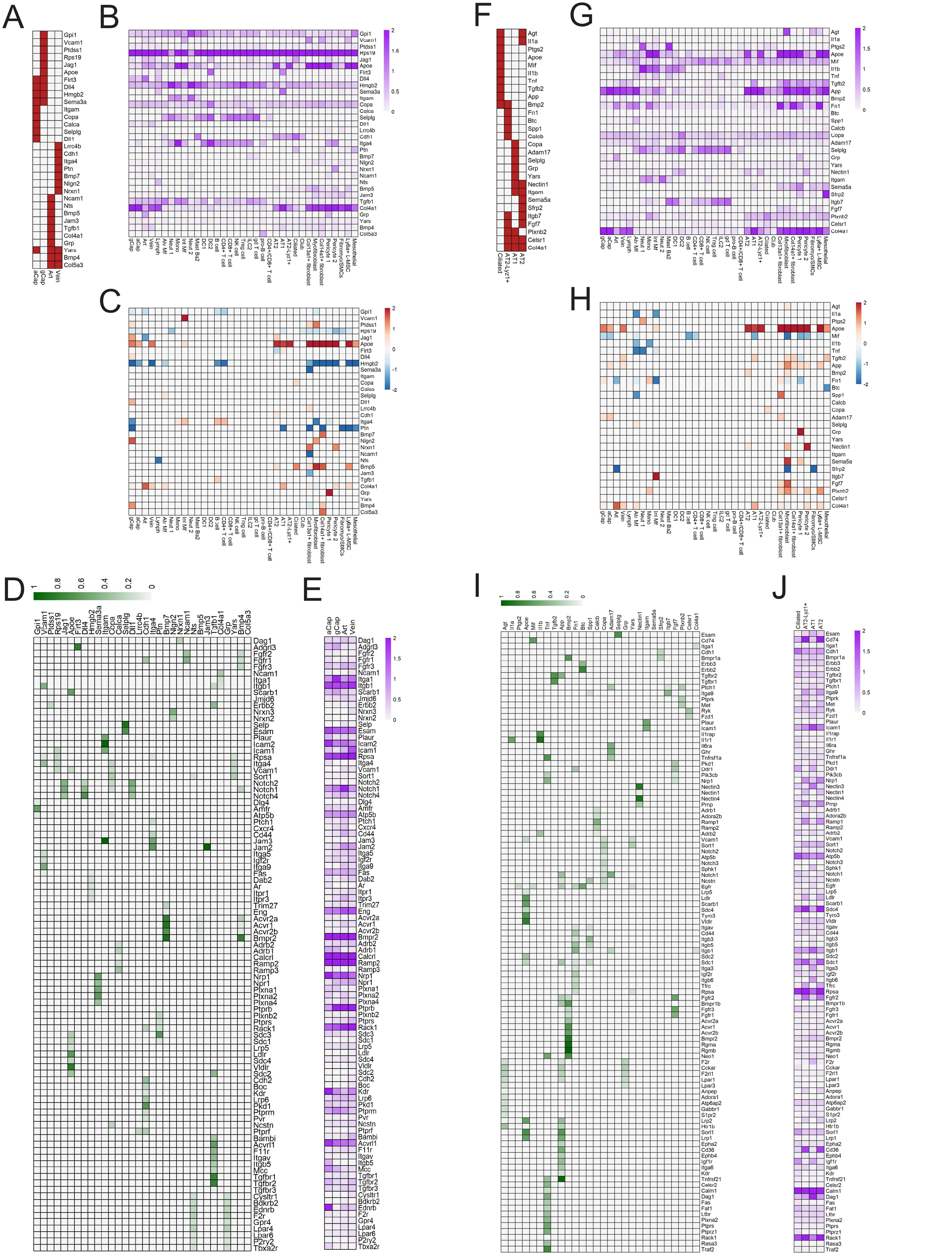


Supplementary Figure 3

### Supplementary Figure 4

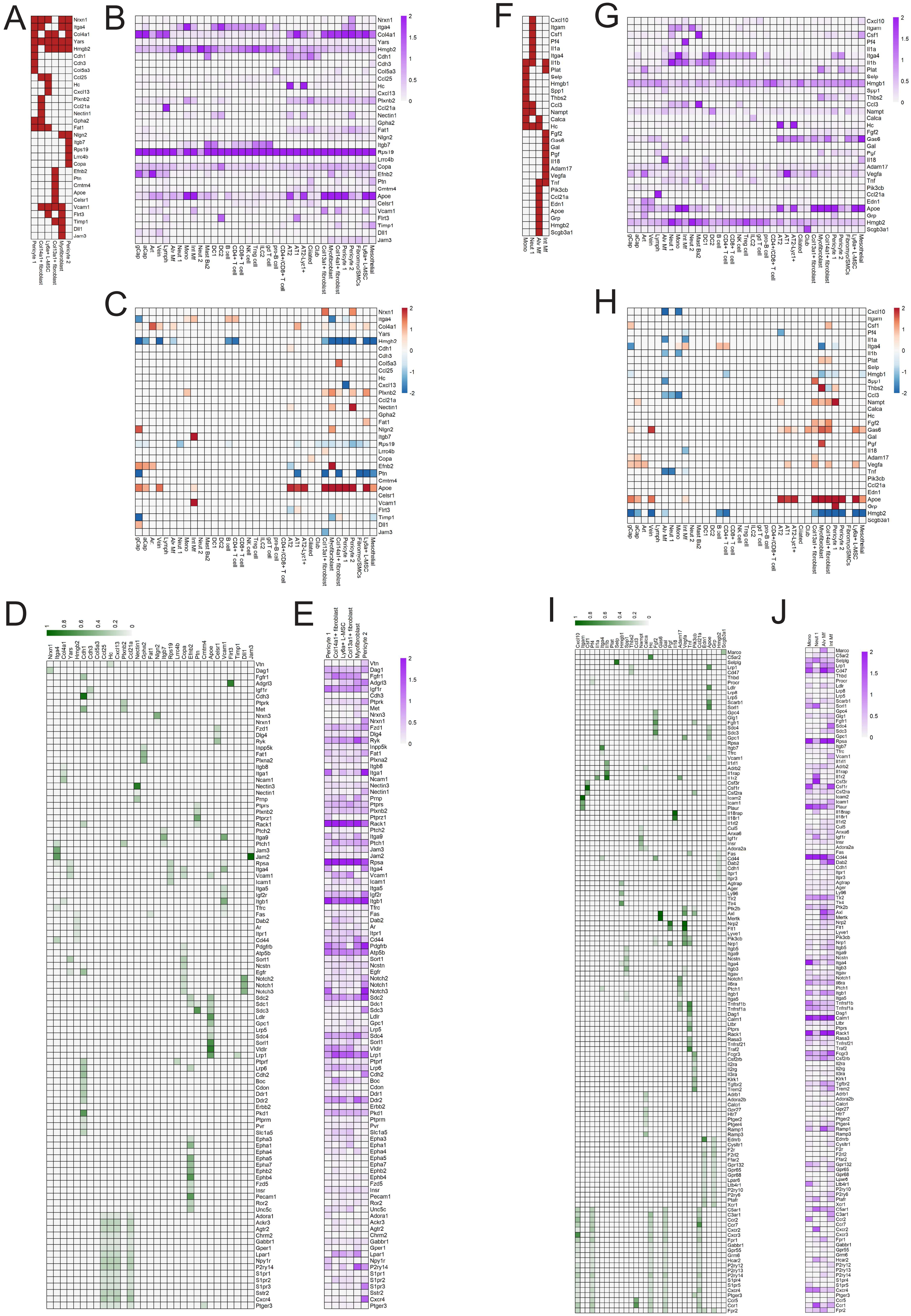


Supplementary Figure 4

### Supplementary Figure 5

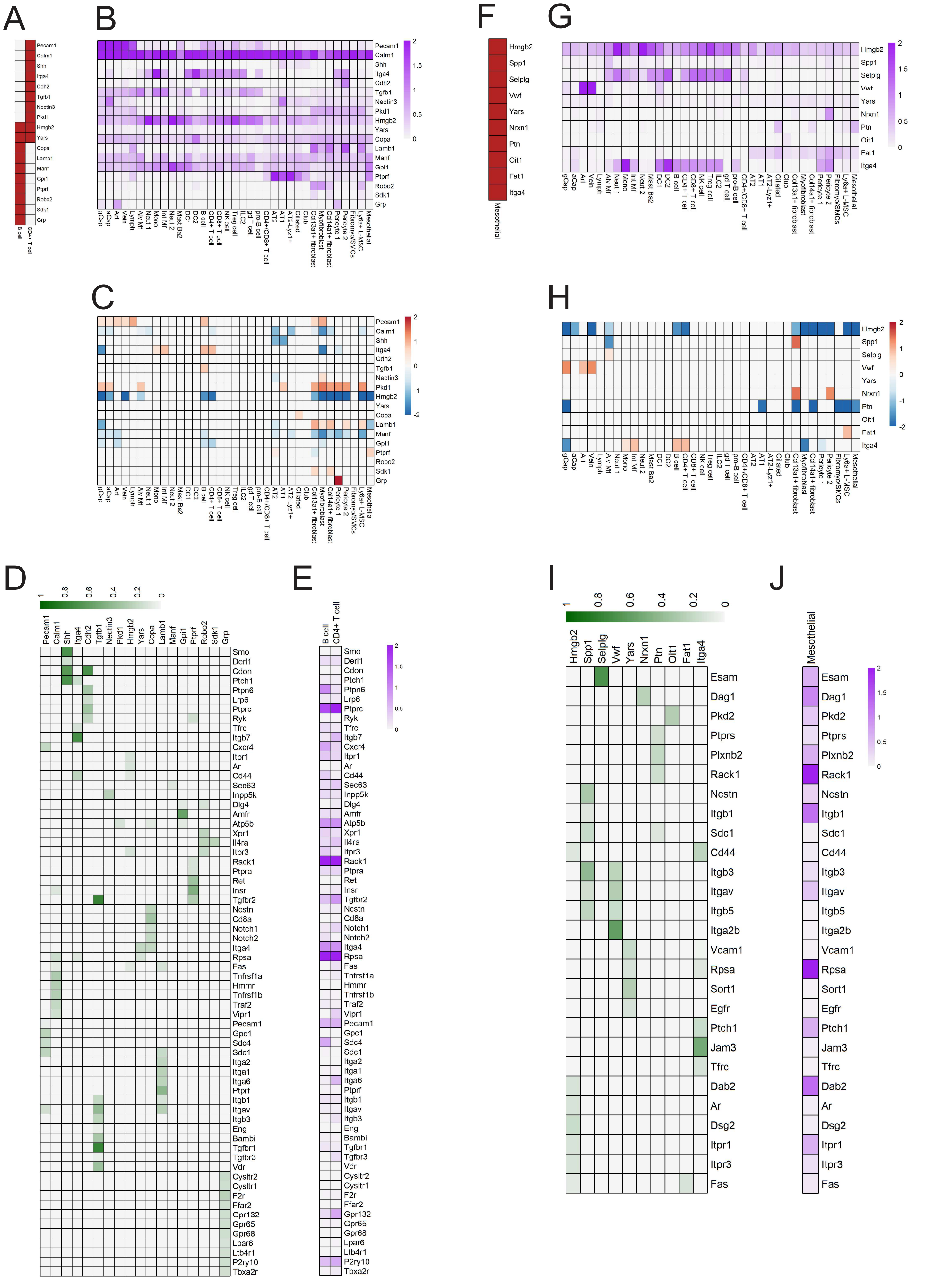


#### Supplementary Figure 5

### Supplementary Figure 6

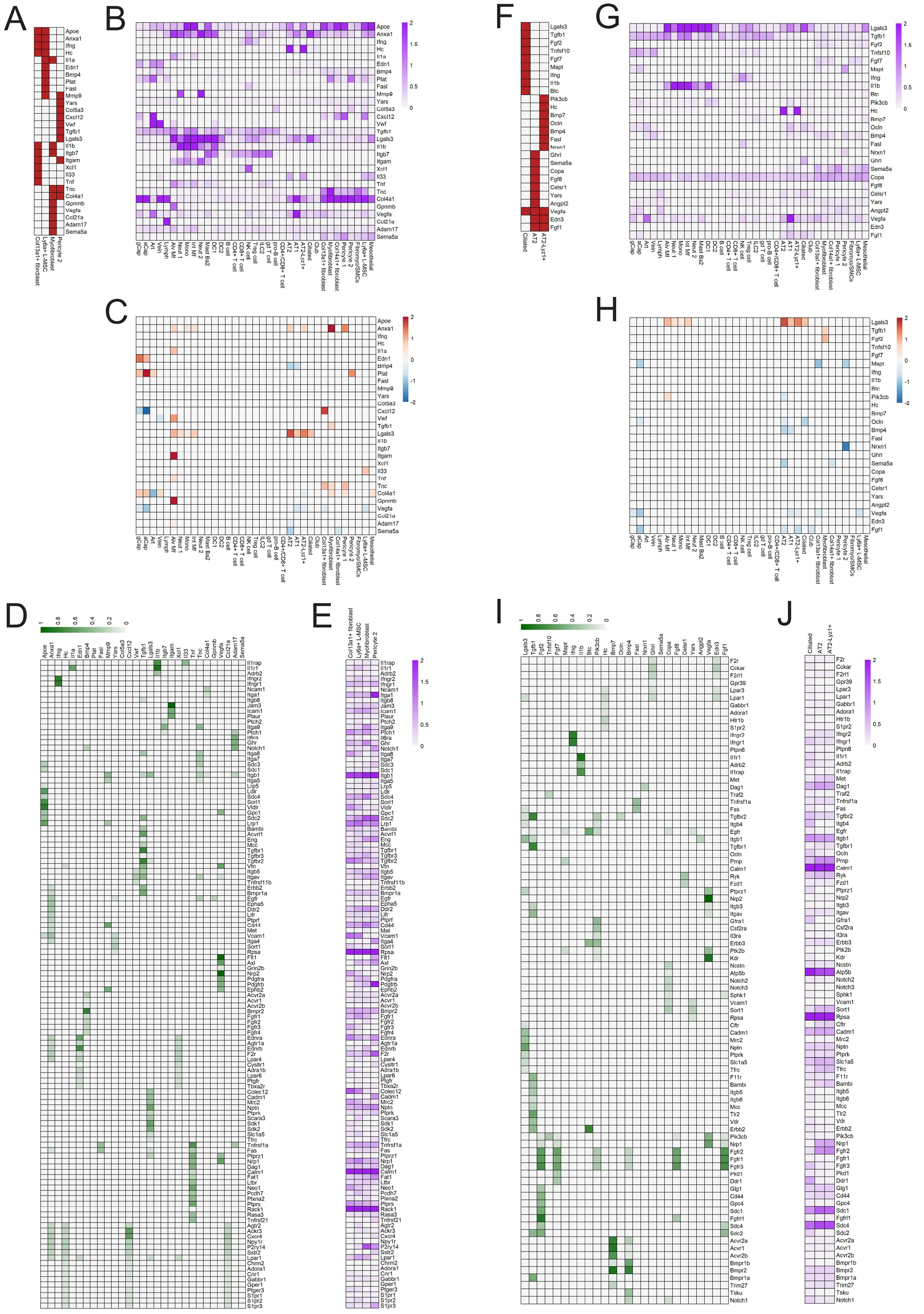


Supplementary Figure 6

### Supplementary Figure 7

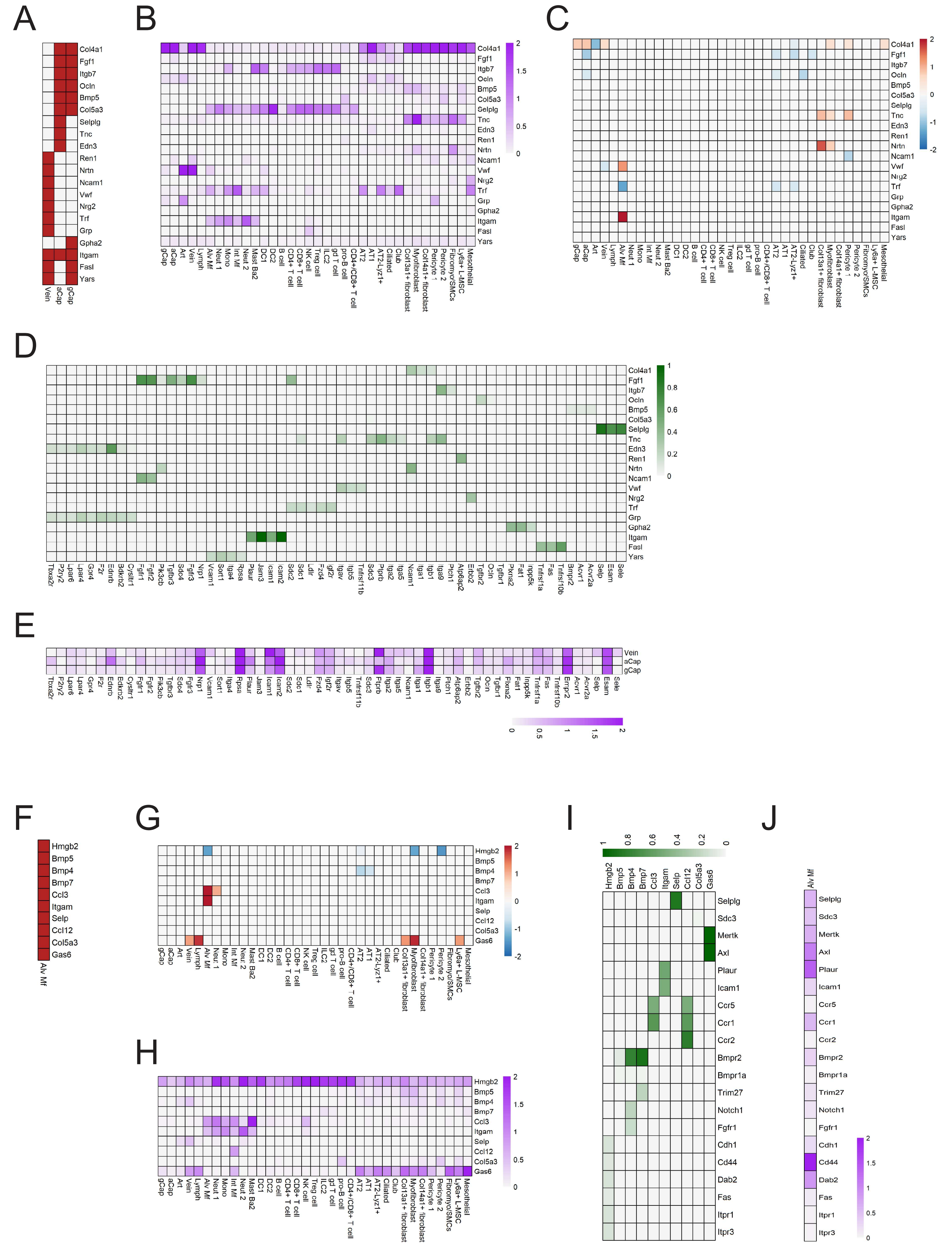


Supplementary Figure 7
