## Supplementary Figures Legends for "Single-cell RNA sequencing-based characterization of resident lung mesenchymal stromal cells in bronchopulmonary dysplasia"

**Supplementary figure 1. Flowchart depicting the allocation of mice to experimental groups.** Flowchart illustrating the group identity of mice sacrificed for the purpuses of the present study. Purple color depicts the groups allocated to hyperoxia (85% $O_2$). Mice sacrificied as a part of previously published scRNA-seq dataset from developing newborn mice are not included.

**Supplementary figure 2. Identification of *Ly6a*$^{+}$ L-MSCs in the developing lung.**
**(A)** UMAP plots depicting the expression of commonly used MSC markers in lung stromal cells isolated from lungs of room air (21% $O_2$) or hyperoxia (85%$O_2$)-exposed developing mice. **(B)** Metascape functional enrichment analysis for ligands indicated in Fig. 2C**.** Developmental age-associated summary pathways relevant to lung are depicted. **(C)** Quantification of FACS results in lung homogenates. n = 3 animals/group. Data are presented as means ± SD. Significance was evaluated by ordinary one-way ANOVA with Tukey multiple comparisons correction test. *P* values < 0.05 were considered significant and are depicted. LY6A$^{+}$ **(D)** L-MSCs were isolated from developing mice at P7 and identified by flow cytometry as CD45$^{-}$/CD31$^{-}$/CD326(EPCAM)$^{-}$ /LY6A(SCA1)$^{+}$ cells. n = 3 animals/group. **(E)** Selected hyperoxia-regulated signalling pathways specific only to the *Ly6a*$^{+}$ L-MSC cluster identified by gene set enrichment analysis (GSEA). All terms are significantly enriched (adjusted p-value < 0.05). Normalized enrichment scores (NES) values were computed by gene set enrichment analysis on fold change-ranked genes. **(F)** Metascape functional enrichment analysis for ligands indicated in Fig. 3E**.** Hyperoxia-regulated summary pathways relevant to lung are depicted. **(G)** Identifying markers were first established in the P7 *in situ*, or cultured *Ly6a*$^{+}$ and *Ly6a*$^{-}$ lung stromal cells based on Supplementary tables 4, 15, 16, 17 and 18, as well as Fig. 3A and 6E. The expression levels of these identifying markers are depicted here in the *in situ* lung stromal cells from the room air or hyperoxia-exposed mice at P7. The intensity of expression is indicated as specified by the colour legend. **(H)** Identifying markers were first established in the P7 *in situ*, or cultured *Ly6a*$^{+}$ and *Ly6a*$^{-}$ lung stromal cells based on Supplementary tables 4, 15, 16, 17 and 18, as well as Fig. 3A and 6E. The expression levels of these identifying markers are depicted here in the cultured lung stromal cells from the room air or hyperoxia-exposed mice. The intensity of expression is indicated as specified by the colour legend. Expression values in Heatmap represent Z-score-transformed log(TP10k+1) values. Log(TP10k+1) corresponds to log-transformed UMIs per 10k.

**Supplementary figure 3. Developmental age-associated ligand and receptor activity affecting lung endothelial and epithelial populations.** Panels A-E relate to endothelial populations, panels F-J relate to epithelial populations. **(A, F)** Heatmap depicting top 10 ligands predicted to affect the listed lung cell populations (coloured red). **(B, G)** Heatmap depicting average log(TP10k+1) expression values of ligands for each cell population in the P14 samples (depicted in violet). **(C, H)** Heatmap depicting the log(fold change) expression of ligands in the P14 samples (depicted in red/blue). **(D, I)** Heatmap depicting putative receptors for each ligand according to the prior interaction potential in NicheNet's model (depicted in green). **(E, J)** Heatmap depicting average log(TP10k+1) expression values of receptors for each cell population (depicted in violet). Expression values in violin plots represent Z-score-transformed log(TP10k+1) values. Expression levels in UMAP plots and Dotplots are presented as log(TP10k+1) values. Log(TP10k+1) corresponds to log-transformed UMIs per 10k.

**Supplementary figure 4. Developmental age-associated ligand and receptor activity affecting lung stromal and myeloid populations.** Panels A-E relate to stromal populations, panels F-J relate to myeloid populations. **(A, F)** Heatmap depicting top 10 ligands predicted to affect the listed lung cell populations (coloured red). **(B, G)** Heatmap depicting average log(TP10k+1) expression values of ligands for each cell population in the P14 samples (depicted in violet). **(C, H)** Heatmap depicting the log(fold change) expression of ligands in the P14 samples (depicted in red/blue). **(D, I)** Heatmap depicting putative receptors for each ligand according to the prior interaction potential in NicheNet's model (depicted in green). **(E, J)** Heatmap depicting average log(TP10k+1) expression values of receptors for each cell population (depicted in violet). Expression values in violin plots represent Z-score-transformed log(TP10k+1) values. Expression levels in UMAP plots and Dotplots are presented as log(TP10k+1) values. Log(TP10k+1) corresponds to log-transformed UMIs per 10k.

**Supplementary figure 5. Developmental age-associated ligand and receptor activity affecting lung lymphoid and mesothelial populations.** Panels A-E relate to lymphoid populations, panels F-J relate to mesothelial populations. **(A, F)** Heatmap depicting top 10 ligands predicted to affect the listed lung cell populations (coloured red). **(B, G)** Heatmap depicting average log(TP10k+1) expression values of ligands for each cell population in the P14 samples (depicted in violet). **(C,**

**H)** Heatmap depicting the log(fold change) expression of ligands in the P14 samples (depicted in red/blue). **(D, I)** Heatmap depicting putative receptors for each ligand according to the prior interaction potential in NicheNet's model (depicted in green). **(E, J)** Heatmap depicting average log(TP10k+1) expression values of receptors for each cell population (depicted in violet). Expression values in violin plots represent Z-score-transformed log(TP10k+1) values. Expression levels in UMAP plots and Dotplots are presented as log(TP10k+1) values. Log(TP10k+1) corresponds to log-transformed UMIs per 10k.

**Supplementary figure 6. Hyperoxia-induced ligand and receptor activity affecting lung stromal and epithelial populations.** Panels A-E relate to stromal populations, panels F-J relate to epithelial populations. **(A, F)** Heatmap depicting top 10 ligands predicted to affect the listed lung cell populations (coloured red). **(B, G)** Heatmap depicting average log(TP10k+1) expression values of ligands for each cell population in the hyperoxia samples (depicted in violet). **(C, H)** Heatmap depicting the log(fold change) expression of ligands in hyperoxia samples (depicted in red/blue). **(D, I)** Heatmap depicting putative receptors for each ligand according to the prior interaction potential in NicheNet's model (depicted in green). **(E, J)** Heatmap depicting average log(TP10k+1) expression values of receptors for each cell population (depicted in violet). Expression values in violin plots represent Z-score-transformed log(TP10k+1) values. Expression levels in UMAP plots and Dotplots are presented as log(TP10k+1) values. Log(TP10k+1) corresponds to log-transformed UMIs per 10k.

**Supplementary figure 7. Hyperoxia-induced ligand and receptor activity affecting lung endothelial and myeloid populations.** Panels A-E relate to endothelial populations, panels F-J relate to myeloid populations. **(A, F)** Heatmap depicting top 10 ligands predicted to affect the listed lung cell populations (coloured red). **(B, G)** Heatmap depicting average log(TP10k+1) expression values of ligands for each cell population in the hyperoxia samples (depicted in violet). **(C, H)** Heatmap depicting the log(fold change) expression of ligands in hyperoxia samples (depicted in red/blue). **(D, I)** Heatmap depicting putative receptors for each ligand according to the prior interaction potential in NicheNet's model (depicted in green). **(E, J)** Heatmap depicting average log(TP10k+1) expression values of receptors for each cell population (depicted in violet). Expression values in violin plots represent Z-score-transformed log(TP10k+1) values. Expression levels in UMAP plots and Dotplots are presented as log(TP10k+1) values. Log(TP10k+1) corresponds to log-transformed UMIs per 10k.
